# SapM mutation to improve the BCG vaccine: genomic, transcriptomic and preclinical safety characterization

**DOI:** 10.1101/486993

**Authors:** Nele Festjens, Kristof Vandewalle, Erica Houthuys, Evelyn Plets, Dieter Vanderschaeghe, Katlyn Borgers, Annelies Van Hecke, Petra Tiels, Nico Callewaert

## Abstract

The *Mycobacterium bovis* Bacille Calmette Guérin (BCG) vaccine shows variable efficacy in protection against adult tuberculosis (TB). Earlier, we have described a BCG mutant vaccine with a transposon insertion in the gene coding for the secreted acid phosphatase SapM, which led to enhanced long-term survival of vaccinated mice challenged with TB infection. To facilitate development of this mutation as part of a future improved live attenuated TB vaccine, we have now characterized the genome and transcriptome of this sapM::Tn mutant versus parental BCG Pasteur. Furthermore, we show that the sapM::Tn mutant had an equal low pathogenicity as WT BCG upon intravenous administration to immunocompromised SCID mice, passing this important safety test. Subsequently, we investigated the clearance of this improved vaccine strain following vaccination and found a more effective innate immune control over the sapM::Tn vaccine bacteria as compared to WT BCG. This leads to a fast contraction of IFNγ producing Th1 and Tc1 cells after sapM::Tn BCG vaccination. These findings corroborate that a live attenuated vaccine that affords improved long-term survival upon TB infection can be obtained by a mutation that further attenuates BCG. These findings suggest that an analysis of the effectiveness of innate immune control of the vaccine bacteria could be instructive also for other live attenuated TB vaccines that are currently under development, and encourage further studies of SapM mutation as a strategy in developing a more protective live attenuated TB vaccine.

## 1. Introduction

Today, TB remains one of the major causes of infectious disease and death throughout the world [1]. About 10 million people developed TB in 2017, with an estimated 1.3 million deaths.

Although new and promising anti-tubercular drugs are in or approaching the clinic, full control over this devastating disease will necessarily require a good, preventive anti-TB vaccine [2]. M. *bovis* Bacille-Calmette-Guérin (BCG) is the only licensed vaccine today, and already in use for almost 100 years. Although BCG is protective against TB meningitis and disseminated TB in children, its efficacy against pulmonary TB is highly variable and not sufficient for disease control [3–5].

Currently, 13 vaccine candidates are under active development and in the process of clinical testing. Besides two live recombinant candidates (VPM1002 and MTBVAC) which are being tested to replace BCG as a priming vaccine, the candidates being mainly variations of booster vaccines with recurrent key mycobacterial antigens [2,6,7]. Several of these vaccine candidates yield enhanced interferon γ (IFNγ)-producing cellular immunity and enhanced control over bacterial replication in the first weeks after TB challenge in animal models. However, reports that show extended long-term survival of vaccinated animals upon TB challenge are much rarer. In the category of priming vaccines, it has been shown that rBCG30 (BCG Tice overexpressing the *M. tb* 30-kDa major secretory protein antigen 85B)-immunized mice survive longer than BCG-immunized mice following TB challenge [8]. SO2 *(phoP* inactivated in *M. tb* strain MT103), the MTBVAC (containing a double deletion in *phoP* and *fadD26)* prototype, conferred greater efficacy than BCG in a high-dose challenge, long-term survival experiment in guinea pigs, while no difference to WT BCG was observed after low-dose challenge [9]. The recombinant BCG strain AFRO-1 (BCG1331 Δ*ureC*::Ω*pfoA*_G137Q_, Ag85A, Ag85B and TB10.4) affords better protection than BCG in mice following aerosol challenge with *Mycobacterium tuberculosis* (*M.tb*) [10]. Additionally, some prime-boost vaccination strategies demonstrated improved long-term survival of M.tb-challenged guinea pigs versus BCG alone: BCG priming followed by the M72 recombinant protein booster vaccine (fusion of Mtb32 and Mtb39 antigens of *Mtb)* [11] and ID93 (combines four antigensbelonging to families of M.tb proteins associated with virulence (Rv2608, Rv3619, Rv3620) or latency (Rv1813)/GLA-SE)) [12].

Earlier, we have demonstrated how a *sapM* transposon mutant of *M. bovis* BCG, hereafter called *sapM*::Tn BCG, resulted in a TB vaccine candidate that provides improved long-term survival in mice upon both systemic and intratracheal challenge with *M. tb* as compared to the parental BCG Pasteur-derived strain (hereafter called WT BCG) [13]. We suggested that the secreted acid phosphatase SapM may have evolved as an important immunomodulatory protein of Mycobacteria. The importance of SapM for mycobacterial immunomodulation was since confirmed by independent laboratories [14–17]. In the present study, we report on the in-depth genomic and transcriptomic analysis of the *sapM*::Tn BCG mutant, to assess which alterations are induced by the transposon insertion in the *sapM* locus.

One of the major impediments to TB vaccine development is the incomplete understanding of the mechanisms of protective immunity against *M.tb,* so far obstructing rational vaccine development [18]. Research on animal models and ongoing clinical trials may reveal markers that correlate with a vaccine’s protective potential. However, as long as we lack these correlates, vaccine design will be challenging. One general idea that dominated immunology of infectious diseases over the last decades, is the T helper type 1/type2 (Th1/Th2) paradigm. Following this model, Th1 cells would protect the host from intracellular pathogens, like *M.tb.* Indeed, the early appearance of Th1 type CD4^+^ T cells secreting IFNγ is necessary for the orchestration of protective immunity in the infected lung [19]. However, controversy has arisen about whether the quantitative extent of IFNγ production in recall experiments has value as a correlate of vaccine-induced protection against TB. Indeed, several reports show that other anti-tuberculosis CD4 T-cell effector functions are at play [20–24].

In this evolving context, we also report on the safety in immunocompromised mice as well as rapid innate control over the *sapM*::Tn vaccine bacteria, with intriguing impact on the induced adaptive immunity. Our findings suggest that a live attenuated TB vaccine that behaves more like an acute, rapidly immune-controlled infection, rather than the protracted chronic infection caused by the current BCG vaccine, may yield an ! immune status that affords more prolonged control of a subsequent TB infection.

## 2. Results

### 2.1 In depth characterization of the *sapM*::Tn BCG versus WT BCG strain

#### 2.1.1 Whole-genome resequencing

The *M. bovis* BCG Pasteur strain 1721, used to create the *sapM::Tn* BCG mutant strain [13,25], differs from the 1173P2 Pasteur vaccine strain by a K43R point mutation in the *rpsL* gene, conferring streptomycine resistance [26], which is useful to avoid contaminations during the lengthy and involved procedures in ordered transposon mutant library production. To investigate whether there are any additional variants between the WT BCG Pasteur and the *sapM*::Tn BCG mutant, we performed a whole-genome resequencing analysis on both strains (paired end 2 × 150 bp), mapping to the *M. bovis* BCG Pasteur 1173P reference genome, resulting in ± 80x average coverage of mapped reads (Suppl. Fig. 1). We could reconfirm the location of the inserted Himar1 transposon, at the TA dinucleotide 8 bp before the *sapM* start codon (Fig. 1). This was also demonstrated by *de novo* assembly of the sequencing reads from the *sapM*::Tn BCG mutant (data not shown). In addition, we performed a probabilistic variant analysis on both WT BCG and *sapM*::Tn BCG mappings and detected only minor modifications (single amino acid change) with unknown impact. These variants are listed in Table 1 and were verified by Sanger sequencing. All variants versus the Pasteur reference are present in both the parental as well as the *sapM*::Tn strains, with only one exception, i.e. the frameshift mutation in the *sugI* gene, coding for sugar-transport integral membrane protein. This mutation probably arose after the library preparation (in a later passage of the strain), since this frameshift mutation is not present in *sapM*::Tn. The exact function of *sugI* is unknown, however, it shows distant sequence similarity to glucose permease GlcP of *S. coelicolor* and the galactose (GalP) and arabinose (AraE) transporters of *E. coli.* Thus, the system is likely to transport a monosaccharide [27]. It is described to be non-essential in the H37Rv strain [28], which explains the absence of phenotype in WT BCG due to the frameshift mutation. The SNV in *sapM*::Tn BCG SugI also likely arose after library preparation since it is not present in WT BCG. Hence, *sugI* appears to be a highly variable gene in the BCG lineage during *in vitro* cultivation, as the three strains each have a different sequence variant. Interestingly, both the parental as well as the *sapM*::Tn strain contain aframe-shift mutation in the gene coding for FadD26, a key enzyme in the biosynthesis pathway of the phthiocerol dimycocerosate (PDIM) class of virulence lipids. This *fadD26* mutation is interesting from a vaccine engineering point-of-view, as it further attenuates virulence [29,30].

**Figure 1.**
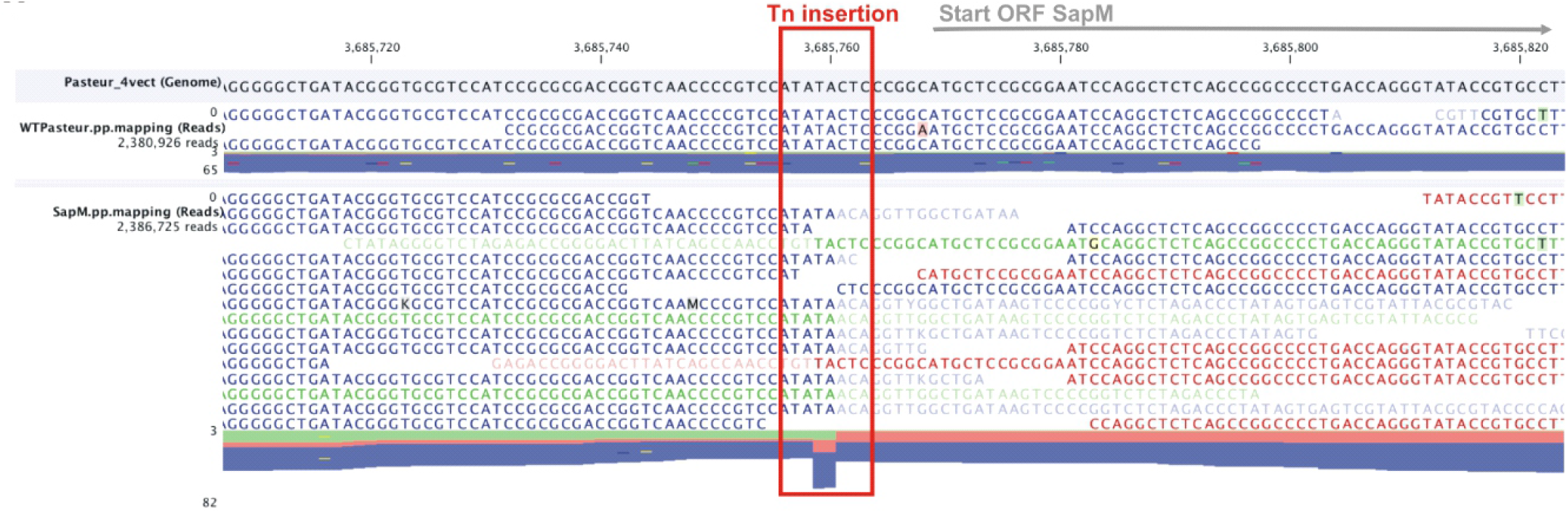
Whole genome resequencing of the WT BCG Pasteur 1721 strain and of the *sapM::Tn* BCG mutant. Detailed representation of the read mappings at the *sapM* locus for both strains. The red square marks the TA dinucleotide site where the transposon is inserted in the mutant strain.

**Table 1.** Variants detected in WT BCG and the *sapM::Tn* BCG disruption mutant. A probabilistic variant analysis was performed in CLC genome workbench on the reads mappings of both the WT BCG and *sapM*::Tn BCG mutant strain. We observed only very few variations compared to the *M. bovis* BCG Pasteur str. 1173P2 reference genome. (SNV/MNV = single/multiple nucleotide variants; Indel = insertions or deletions). The Tn insertion was picked up using *de novo* assembly of the *sapM::Tn* reads.

|  | Position | Mutation Type | Gene | AA change | Illumina resequencing |  |  | Sanger analysis |  |  |
| --- | --- | --- | --- | --- | --- | --- | --- | --- | --- | --- |
|  |  |  |  |  | 1173P2 WT (reference) | 1721 BCG WT | 1721 *SapM* ::Tn | 1173P2 WT | 1721 BCG WT | 1721 *SapM* ::Tn |
| Variant 1 | 813,096 | SNV | *rpsL* | K43R | A | G | G | A | G | G |
| Variant 2 | 1,344,671 | MNV | - | - | CG | GC | GC | GC | GC | GC |
|  | 2,764,157 | SNV* | BCG_2507c | Synonymous | T | A | A |  |  |  |
| Variant 3 | 3,197,940 | InDel | *fadD26* | L85fs | - | A | A | - | A | A |
| Variant 4 | 3,833,491 | InDel** | BCG_3499c | A433InsA | - | CGC | CGC |  |  |  |
| Variant 5 | 3,854,971 | InDel | *cut3* | G261Deletion<br>Insertion GR | - | TCG | TCG | TCG | TCG | TCG |
|  | 3,685,759 | Tn insertion | SapM promoter | - | - | - | tn | - |  |  |
| Variant 6a | 3,708,425 | InDel | *sugI* | S13fs | - | A | - | - | A | - |
| Variant 6b | 3,708,517 | SNV | *sugI* | L44M | C | C | A | C | C | A |

\* Not verified by Sanger sequencing, because the mutation is synonymous
\*\* Sequencing did not work, due to the presence of repeats in the region of the mutation

#### 2.1.2 Transposon-induced transcriptional effects

Transcriptome analysis, by both RNAseq analysis and RT-PCR, demonstrated that the *sapM* transcript is ~20-50-fold reduced compared to the WT BCG, while the transcription levels of other genes remain largely unchanged (< 2-fold change), except for a ~10-fold upregulation of *upp* (encoding uracil phosphoribosyltransferase) which is immediately upstream of *sapM* in the genome, and a modest 2.4-fold downregulation of its downstream gene *(BCG_3376)* (Fig. 2A-C, Suppl. Fig. 2A-B, Suppl. Table 1). A few (~6) other genes show a differential expression pattern (~2-fold change) between WT BCG and *sapM*::Tn, however, this effect is marginal compared to the differences in expression of *sapM* and *upp.* The transposon insertion thus only majorly affects one nearby gene, and the changes in *sapM* and *upp* transcription have almost no secondary effects on overall transcriptional regulation. According to the Operon Correlation Browser on the TB Database [31], the gene coding for SapM in *M. tb* (Rv3310) and the downstream gene Rv3311 potentially form an operon. However, in *M. bovis* BCG we see that transposon disruption of the *sapM* upstream region in *sapM*::Tn BCG very strongly reduces the levels of *sapM* transcript, while having a much more modest effect on the levels of BCG_3376 (Rv3311 ortholog) transcription. These data indicate that both genes are likely differentially regulated, questioning the idea of an operonic structure.

**Figure 2.**
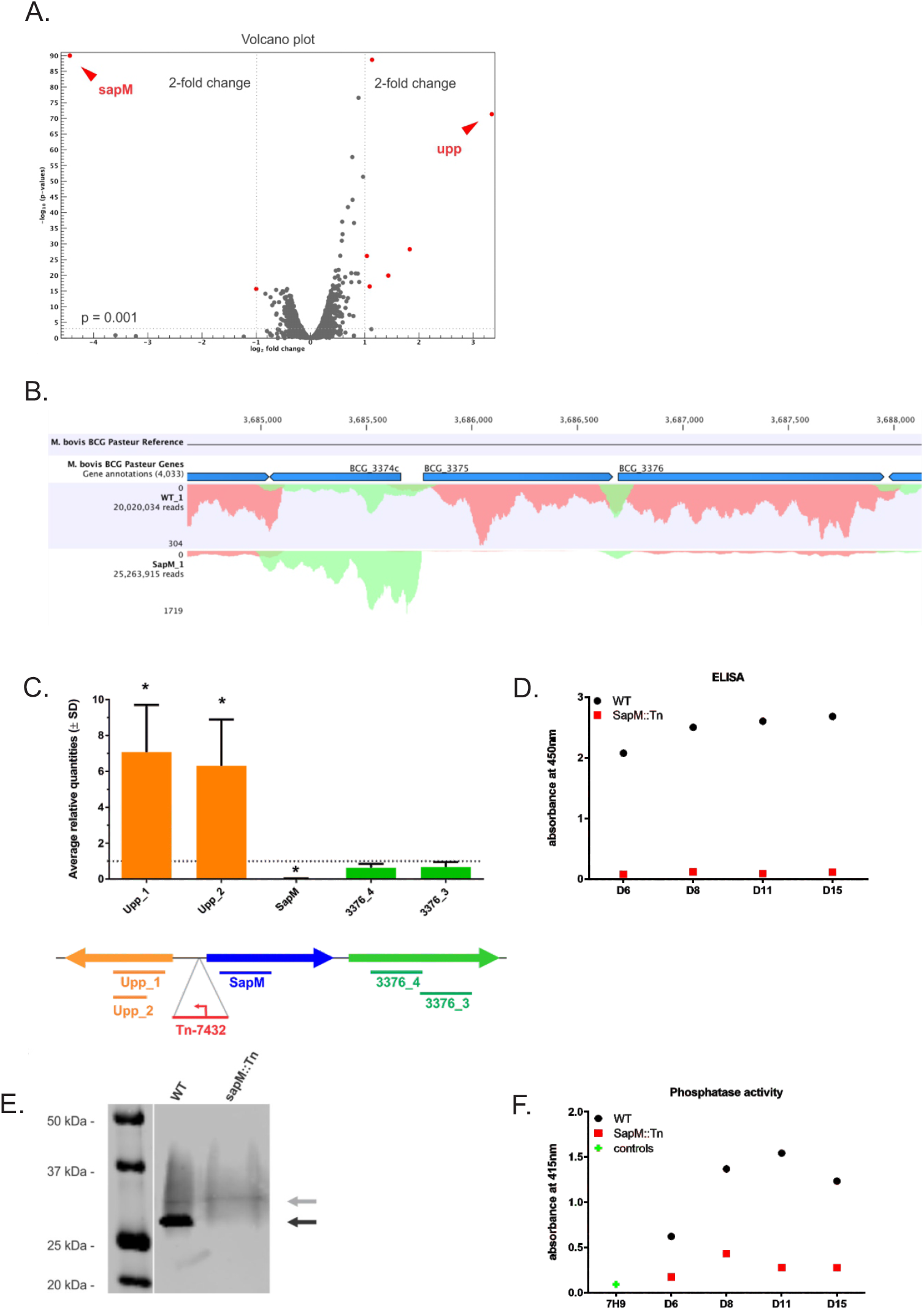
(A) Volcano plot comparing the *sapM*::Tn BCG mutant and the WT BCG. Genes with a > 2-fold change and a p-value < 0.001 are depicted in red. **(B) Mapped (sense) reads at the SapM locus in the WT BCG vs *sapM*::Tn BCG condition**. Only one replicate of both WT BCG and *sapM*::Tn BCG samples is depicted. BCG_3374c = *upp,* BCG_3375 = *sapM.* **(C) RT-PCR analysis of the *sapM* locus**. RNA was prepared of cultures of biological triplicates of the *sapM*::Tn BCG disruption mutant, as well as of the WT BCG. An RT-PCR on the cDNA using primer sets directed against the sapM gene (blue bar) and the directly up- and downstream genes *(upp* and *BCG_3376,* orange and green bars, respectively). The data presented here are averages (± SD) of the three biological replicates. For each mutant, the targets are individually normalized to the transcription levels in the WT BCG strain (grey dotted line set at an average relative quantity of 1). The * indicates significance (p < 0.05) by a two tailed t-test comparing the transcription levels of each target in the mutant strain to the same target in the WT BCG. (D-F) **SapM protein and activity analysis**. The *sapM*::Tn BCG mutant and WT BCG 1721 were grown in 7H9 medium and supernatant samples were collected on various time points (D6 (Day 6) – D15). **(D)** ELISA using an anti-SapM polyclonal antibody. **(E)** SDS-PAGE and western blotting of WT BCG and *sapM*::Tn BCG mutant supernatant samples collected on day 15 and grown in 7H9 medium without OADC and supplemented with glucose only. The blots were developed with the anti-SapM antibody. The processed SapM protein (= removal of the N-terminal signal sequence) is ± 28 kDa in size and is only present in the WT BCG strain (black arrow). **(F)** *In vitro* phosphatase assay using p-Nitrophenyl phosphate (pNPP) as a substrate to check the activity of the SapM enzyme. The plotted data for both ELISA and phosphatase assay are averaged over 2 technical replicates and corrected for the background signal induced by sterile 7H9 medium supplemented with 10% OADC.

#### 2.1.3 SapM protein is undetectable and secreted total phosphatase activity is strongly reduced

As the SapM protein contains a 43 AA N-terminal signal sequence and thus gets secreted in the culture medium, we collected culture supernatant samples at various time-points.

First, we performed an ELISA on these supernatant samples with a newly generated antibody directed against the recombinant SapM protein produced in *E. coli* (Fig. 2D). While the amount of SapM protein steadily increases over time of culture in the WT BCG, the signal is nearly absent in the *sapM::Tn* BCG mutant samples for all time-points. We further confirmed the absence of full-length SapM protein in the *sapM*::Tn BCG strain by SDS-PAGE and western blotting in an independent experiment (Fig. 2E). We detected a band of ± 28 kDa in the WT BCG, but not in the *sapM*::Tn BCG mutant sample. A smear is also detected above the 28 kDa band, which likely represents aspecific binding (as the pattern appears to be identical for all samples). To check for any remaining phosphatase activity in the culture medium, we performed an *in vitro* phosphatase assay with p-Nitrophenyl phosphate (pNPP) as a substrate [32] (Fig. 2F). The measured *in vitro* activity is significantly reduced for the *sapM*::Tn BCG mutant sample compared to WT BCG at early time-points. At later time-points (day 8), we still observe a considerable signal, which is expected, as the assay would also pick up activity of other phosphatases such as the protein tyrosine phosphatases (PtpA and PtpB) [33].

Consequently, based on all of the data on this vaccine strain, we concluded at this point that its improved vaccine efficiency must indeed be due to a loss of function of SapM or the upregulation of *upp* transcription, or both.

### 2.2 *In vivo* safety of *sapM*::Tn BCG versus WT BCG

A requirement of live attenuated vaccines is that they should be safe, even in immunocompromised hosts. Our previous analysis of bacterial replication in immunocompetent Balb/c mice, infected intravenously, did not show any difference between WT BCG and *sapM*::Tn BCG [13]. The number of granulomas formed in livers and lungs post-infection was unaffected [13]. To unambiguously demonstrate safety of the *sapM*::Tn BCG mutant compared to WT BCG *in vivo,* we investigated bacterial virulence in the absence of adaptive immunity. Immunocompromised SCID mice, lacking both T and B cells, were infected intravenously (i.v.) with a high (3 × 10^7^ CFU) or lower dose (3 × 10^6^ CFU) of both strains and survival was monitored (Fig. 3). After both low and high doses, SCID mice infected with *sapM*::Tn BCG mutant showed similar survival to mice infected with WT BCG. Thus, in the SCID model, virulence of *sapM::Tn* BCG is comparable to WT BCG.

**Figure 3.**
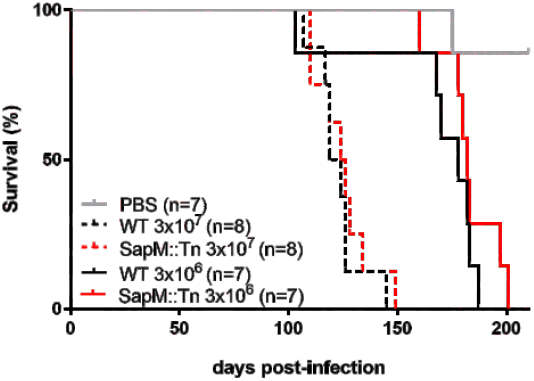
SapM::Tn BCG safety study in SCID mice. Immunocompromised SCID mice were infected intravenously with 3 × 10^6^ versus 3 × 10^7^ cfu of the WT BCG or SapM::Tn BCG mutant and survival was monitored. (Log-rank test; WT BCG vs. *sapM::Tn;* P=not significant). The control groups represent animals vaccinated with PBS.

### 2.3 At the lymph node draining the vaccination site, the *sapM*::Tn BCG mutant is controlled faster than the WT BCG strain

Our previously reported observations indicated that vaccination with the *sapM*::Tn BCG mutant triggered more effective recruitment of iDCs into the draining lymph nodes (LNs) in Balb/c mice [13]. This lead us to study the influence of this more effective induction of the innate immune response at early time points after *sapM*::Tn BCG vaccination on the fate of the vaccine inoculum. To enhance the validity, experiments were performed in F1 mice (C57BL/6J x Balb/c) to assess robustness of the observed effects to a more heterogeneous immunogenetic background [34]. After subcutaneous (s.c) vaccination of F1 mice, less bacteria (~50%) were recovered from LNs of *sapM*::Tn BCG infected mice as compared to WT BCG-infected mice at all time-points analyzed (i.e. days 8, 14 and 28 post-infection) (Fig. 4A). It is clear that whereas WT BCG could amplify from the inoculum, this was barely the case for the *sapM*::Tn BCG mutant. Since differences are clear already 8 days post-vaccination, this result is most consistent with improved control by innate immune mechanisms. Indeed, an analysis of intracellular survival of WT BCG and *sapM*::Tn BCG upon infection of bone marrow derived macrophages (BM-DMs), showed that WT BCG and *sapM*::Tn BCG are taken up similarly (as shown before [13]) but that growth of the *sapM*::Tn BCG mutant is better controlled compared to the WT BCG strain upon infection (Fig. 4B). In line with these findings, such improved clearance of SapM-mutated *M.tb* by BM-DMs has recently also been described [16], an effect that was attributed to improved phagosomal maturation.

**Figure 4.**
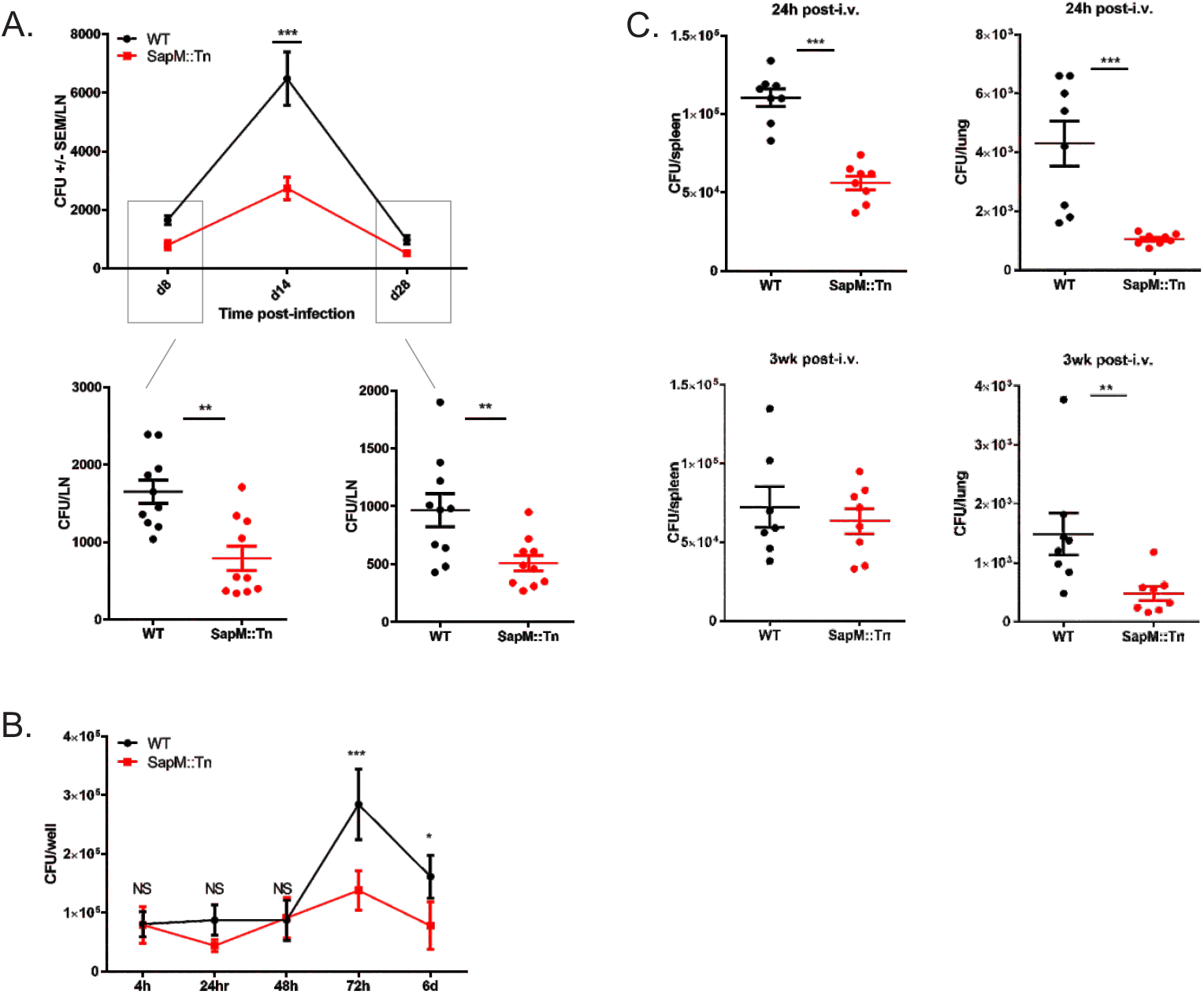
*In vivo* and *in vitro* replication analysis: *sapM*::Tn BCG versus WT BCG. (A) F1 mice (Balb/c x C57BL/6J) were vaccinated s.c. with the WT BCG or *sapM*::Tn BCG (2 × 10^6^ cfu, 10 mice/group). At day 8, 14 and 28 post-infection, mice were sacrificed and the number of bacteria in the draining LNs was determined by cfu plating (Mann-Whitney test; ** P<0.01). **(B)** BM-DMs were infected with WT BCG or *sapM*::Tn BCG (MOI 10:1) for 4 hours. Cells were washed and bacterial uptake was determined 4hr post-infection. Bacterial replication was analyzed by cfu plating 24, 48 and 72hr, 6 days post-infection. (C) F1 mice were vaccinated i.v. with the WT BCG or *sapM*::Tn BCG (2 × 10^6^ cfu, 8 mice/group). At 24h and 3 weeks post-infection, mice were sacrificed and the number of bacteria in the spleens and lungs was determined by cfu plating (Mann-Whitney test; ** P<0.01).

To further evaluate the impact of early innate growth control of the bacteria, we have analyzed survival of *sapM*::Tn BCG and WT BCG after intravenous infection of F1 mice. Twenty-four hours post-infection, the bacterial load in the lungs and spleens from mice that were infected with WT BCG was higher than the loads of those infected with the *sapM*::Tn BCG mutant (~50%) (Fig. 4C). Three weeks post-infection, the difference was still observed in the lungs, but not in the spleens. Similar results were obtained in the spleen of C57BL/6J mice, but not in the lungs (Suppl. Fig. 3A). Cytokine levels were analyzed in the serum of i.v. exposed mice at different time-points (6h, 24h, 48h post-vaccination), and significantly higher levels of IL1-β and IL17 could be measured 24h post-*sapM*::Tn BCG vaccination compared to WT BCG vaccination (Suppl. Fig. 3B). In view of this early time point post-vaccination, both IL1-β and IL17 are produced by innate immune cells. These particular cytokines are known to be important innate cytokines for bacterial killing, particularly in mycobacterial infection [35,36], which is consistent with the reduced *sapM*::Tn BCG bacterial load.

### 2.4 Complementation of the *sapM*::Tn mutation reverses the phenotypes

As the Tn insertion in the *sapM* locus caused both a loss of function of *sapM* and a strong upregulation of the upstream *upp* gene, we assessed whether the improved early innate growth control of the *sapM*::Tn BCG bacteria was due to the loss of function of SapM or the upregulation of *upp* transcription. For this purpose, we have complemented the *sapM*::Tn BCG mutant strain with the *sapM* gene under control of its own promotor. The *sapM* expression is reverted to wild type levels in this *sapM*::Tn:compl BCG mutant, but *upp* levels are still increased similarly as in the *sapM*::Tn BCG mutant (Fig. 5). Analysis of bacterial load in the lungs and spleens from mice that were infected with either *sapM*::Tn:compl BCG mutant, *sapM*::Tn BCG or WT BCG showed that complementation of *sapM* expression abolished the improved growth control of the *sapM*::Tn BCG mutant compared to the WT BCG strain upon infection (Fig. 6). These results confirm that the observed phenotypes are due to the reduced expression of *sapM* and not because of upregulation of *upp* (Fig. 2C).

**Figure 5.**
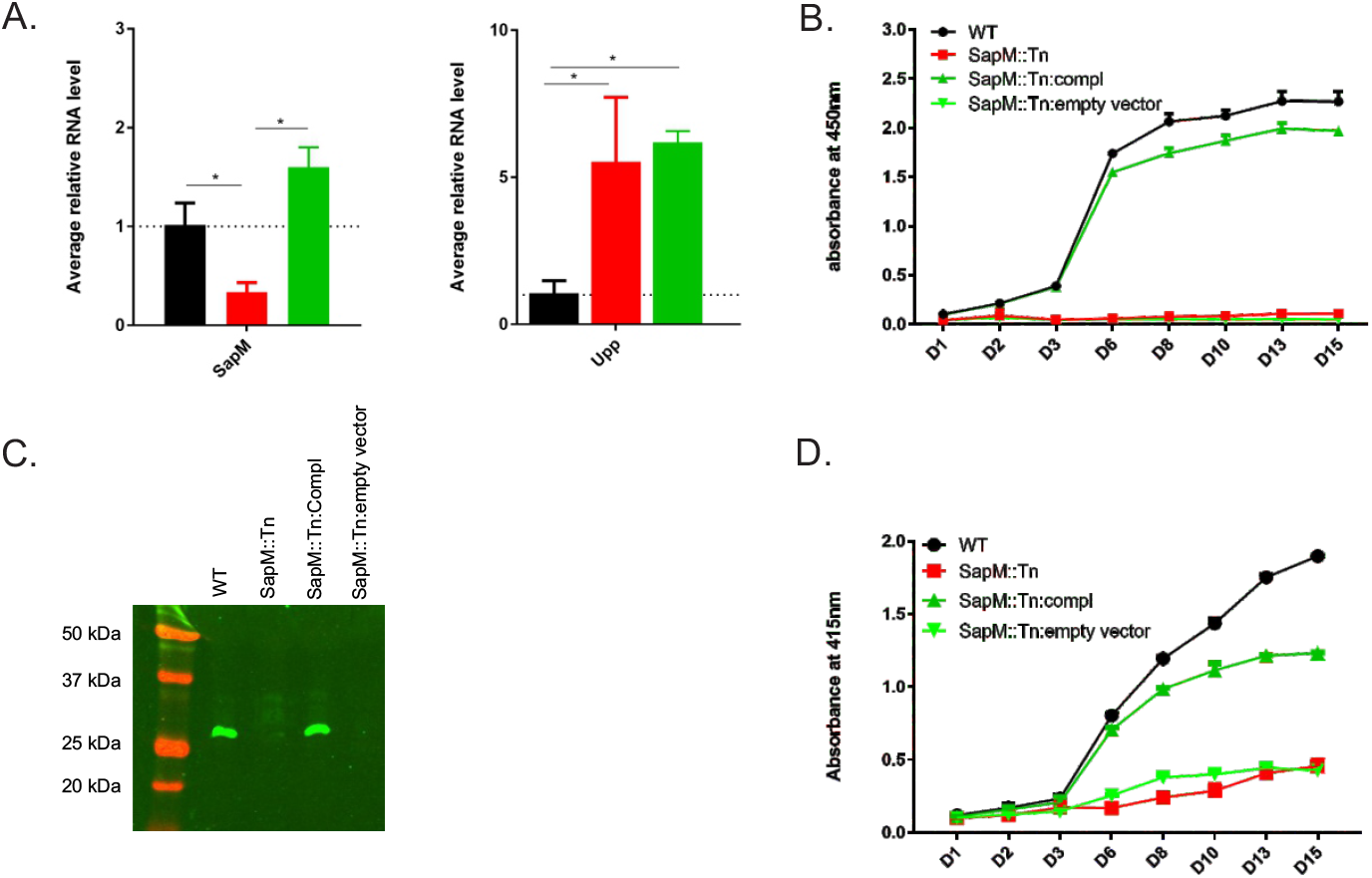
RT-PCR analysis of the *sapM* locus and SapM protein and activity analysis in the *sapM*::Tn:complementation mutant versus WT BCG. **(A)** RNA was prepared of cultures of biological triplicates of the *sapM*::Tn BCG disruption mutant (red), the WT BCG (black) and the *sapM*::Tn:complementation BCG mutant (green). An RT-PCR on the cDNA used primer sets directed against the sapM gene (sapM b primers – left panel) and the directly upstream gene upp (upp2 primers – right panel). The data presented here are averages (± SD) of three technical replicates. For each mutant, the targets are individually normalized to the transcription levels in the WT BCG strain (grey dotted line set at an average relative quantity of 1). The * indicates significance (p < 0.05) by a t-test comparing the transcription levels of each target in the mutant strain to the same target in the WT BCG. **(B-D)** The *sapM*::Tn BCG mutant, WT BCG 1721, *sapM*::Tn:complementation BCG mutant and *sapM*::Tn:empty vector were grown in 7H9 medium and supernatant samples were collected on various time points (D1 (Day 1) – D15). **(B)** ELISA using an anti-SapM polyclonal antibody. The plotted data for both ELISA and phosphatase assay is averaged over 2 technical replicates and corrected for the background signal induced by sterile 7H9 medium supplemented with 10% OADC. (C) SDS-PAGE and western blotting of indicated supernatant samples collected on day 10 and grown in 7H9 medium without OADC and supplemented with glucose only. The blots were developed with the anti-SapM antibody. The processed SapM protein (= removal of the N-terminal signal sequence) is ± 28 kDa in size and is only present in the WT BCG strain and *sapM*::Tn:complementation mutant. (D) *In vitro* phosphatase assay using p-Nitrophenyl phosphate (pNPP) as a substrate to check the activity of the SapM enzyme.

**Figure 6.**
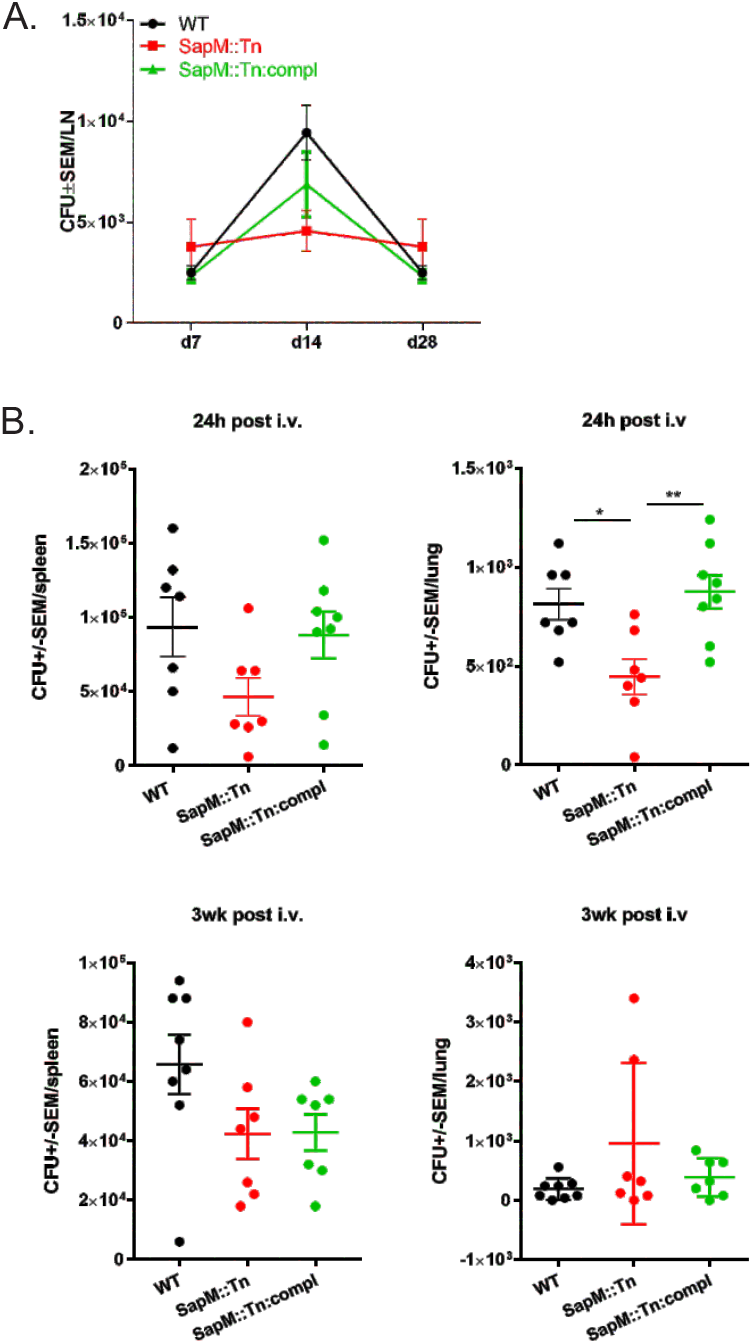
*In vivo* replication analysis: *sapM*::Tn BCG versus *sapM*::Tn:complementation BCG versus WT BCG. **(A)** C57BL/6J mice were vaccinated s.c. with the WT BCG, *sapM*::Tn BCG or *sapM*::Tn:complementation BCG (2 × 10^6^ cfu, 6 mice/group). At day 8, 14 and 28 post-infection, mice were sacrificed and the number of bacteria in the draining LNs was determined by cfu plating (Mann-Whitney test; ** P<0.01). **(B)** C57BL/6J mice were vaccinated i.v. with the WT BCG, *sapM*::Tn BCG or *sapM*::Tn:complementation BCG (2 × 10^6^ cfu, 7-8 mice/group). At 24h and 3 weeks post-infection, mice were sacrificed and the number of bacteria in the spleens and lungs was determined by cfu plating (Mann-Whitney test; ** P<0.01).

### 2.5 Faster kinetics of iDC recruitment to lymphoid organs and decrease in IFNγ-producing CD4^+^ and CD8^+^ T recall response in *sapM*::Tn BCG vaccinated mice compared to WT BCG vaccinated mice

We then further analyzed how improved innate control over vaccine bacteria changes the immune response to the vaccine. Hereto, we have vaccinated C57BL/6J mice s.c. with WT BCG or *sapM*::Tn BCG mutant. At different time points, mice were sacrificed and LNs and spleens were isolated. DC phenotyping was performed. Significant differences were observed in total leukocyte numbers and iDC recruitment kinetics after *sapM*::Tn BCG versus WT BCG vaccination. In general, we observe a faster induction of the innate immune response, reflected by a faster recruitment of innate inflammatory cells to the secondary lymphoid organs (dLN-spleen), and by a faster contraction of the innate immune response, seen by lower leukocyte cell numbers at day 14 post-vaccination with BCG *sapM*::Tn BCG (Fig. 7). Next, T cell immunity was assessed by restimulation of splenocytes or LN cells with either the mycobacterial antigen Ag85A, the *M.tb* extract PPD or by anti-CD3/28, a stimulus that activates all T cells, as a control. Vaccination with *sapM*::Tn BCG leads to a lower number of lymph node and spleen IFNγ-producing CD4^+^ (Th1) and CD8^+^ (Tc1) T cells 2 weeks postvaccination compared to WT BCG (Fig. 8A). After vaccination with a lower dose, this observation is only seen for Tc1 cells, following polyclonal stimulation in the LNs and spleen (Suppl. Fig. 4). IFNγ measurements in the supernatant following stimulation with the different antigens also demonstrated lower levels of IFNγ in the *sapM*::Tn BCG condition (Fig. 8B). These lower numbers of IFNγ-producing CD4^+^ and CD8^+^ T cells were seen at early time points (2 weeks post-infection), but the difference became non-significant at later time points (Suppl. Fig. 5), consistent with the hypothesis that a fast innate control of bacterial load results in a swift contraction of the TB-specific Th1 and Tc1 cellular response after *sapM*::Tn BCG vaccination. The *sapM*::Tn BCG vaccine thus behaves like a rapidly (though not completely) controlled bacterial infection, contrary to WT BCG, which expands and causes a more protracted infection at the vaccination site.

**Figure 7.**
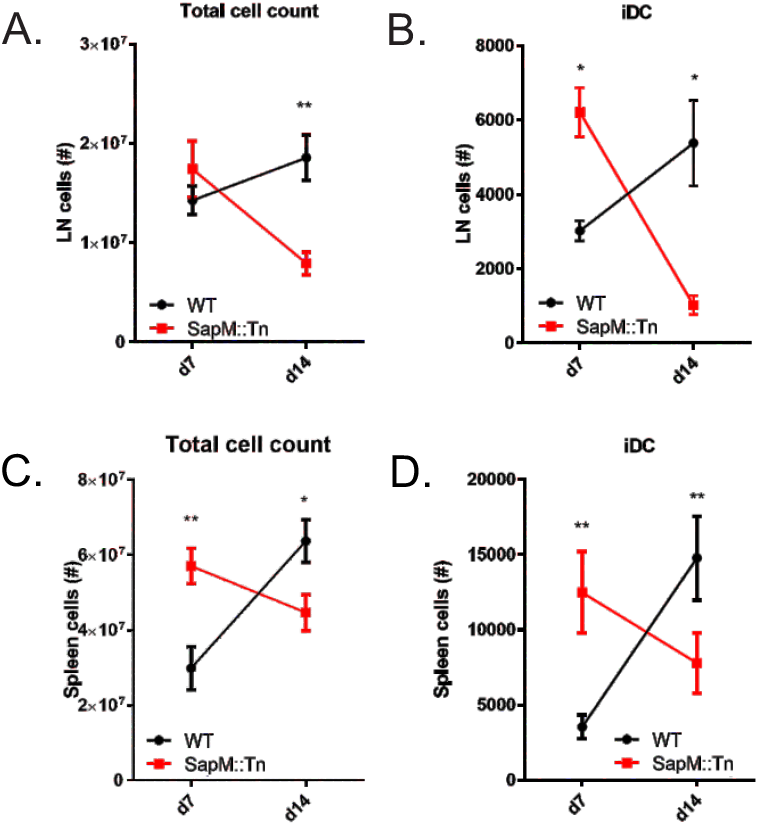
iDC recruitment to lymphoid organs starts earlier when mice are vaccinated with the BCG *sapM*::Tn BCG strain. C57BL/6J mice were immunized s.c. at the base of the tail (2 × 10^6^ cfu, 5-10 mice/group (A-B); 2.10^5^ cfu, 10 mice/group (C-D)) with WT BCG or *sapM*::Tn BCG mutant. Seven and 14 days later, the mice were sacrificed, and inguinal LNs, brachial LNs and spleens were isolated. Cells were prepared, labelled with different antibodies staining DCs and analyzed by flow cytometry. Total cell numbers **(A, C)** and the kinetics of iDC (CD103+MHCII^hl^ CD11b+ Ly6C+) recruitment **(B, D)** differ upon BCG *sapM*::Tn BCG and WT BCG vaccination. (Mann-Whitney; *P<0.05, **P<0.01).

**Figure 8.**
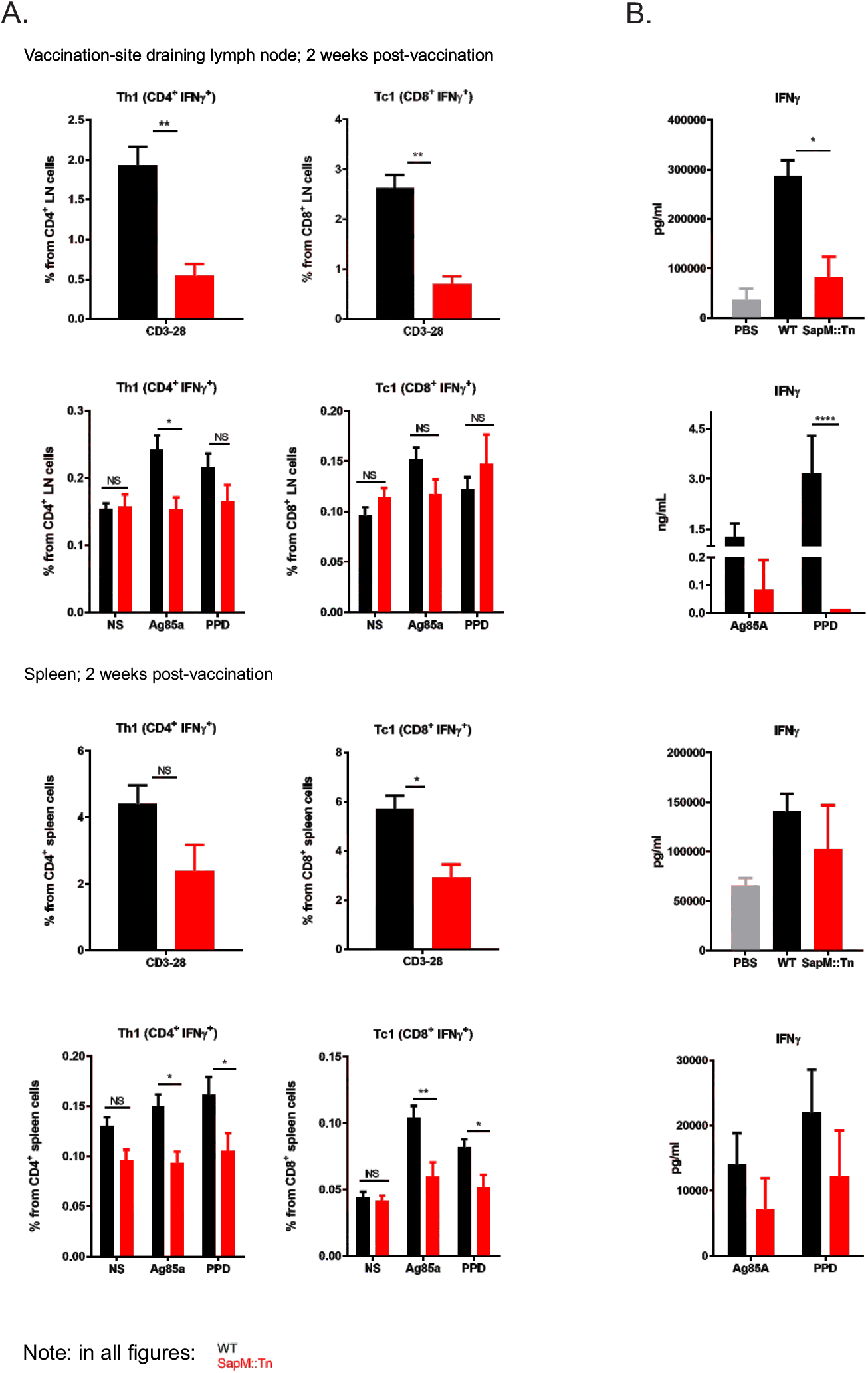
Vaccination with *sapM*::Tn BCG induces reduced frequencies of IFNγ-producing CD4^+^ and CD8^+^ T cells compared to WT BCG. C57BL/6J mice were immunized s.c. at the base of the tail with a high dose of WT BCG (black bars) or *sapM*::Tn BCG mutant (red bars) (2 × 10^6^ cfu, 5 mice/group). 14 days later, the mice were sacrificed, and inguinal LNs, brachial LNs and spleens were isolated. Cells were prepared and the T cell response was analyzed by intracellular cytokine staining followed by flow cytometry **(A)**. Additionally, supernatants of stimulated spleen cells and controls were harvested 48hr post-stimulation and IFNγ concentration was determined by bioplex **(B)**. (Mann-Whitney or 2way ANOVA; *P<0.05, **P<0.01).

## 3. Discussion

Our former research demonstrated that an *M. bovis* BCG *sapM*::Tn mutant led to enhanced long-term survival of vaccinated mice challenged with TB [13]. Further development of this TB vaccine candidate requires further characterization of the mutant and its safety, which was the main purpose of the extensive work presented in this study.

We have performed a whole genome resequencing-based variant analysis of this *sapM*::Tn BCG mutant, as well as of its parent BCG strain [26], using a shotgun Illumina sequencing approach. Very few polymorphisms could be detected compared to the *M. bovis* BCG Pasteur reference genome, most of which were already present in the streptomycin-resistant Pasteur derivative in which we designed our transposon mutant. A frame-shift mutation had inactivated the gene coding for FadD26, a key enzyme in the biosynthesis pathway of the PDIM class of virulence lipids that are involved in hiding *M. tb’s* own PAMPs from the host’s innate immune system [29]. Knocking out this gene in *M. tb* severely impairs the pathogen’s ability to survive *in vivo* [29,30]. For this reason, the *fadD26* gene is deleted in MTBVAC, the first live-attenuated *M. tb* vaccine [37]. Chen *et al.* compared PDIM/PGL production in 12 different *M. bovis* BCG substrains and found that while most were PDIM-positive (including BCG Pasteur), three of these strains, BCG Japan (or Tokyo), Moreau and Glaxo, do not produce this class of lipids [38]. The authors further reported that there was a correlation between the virulence that was associated with the BCG substrains and their lipid profile: the on average more virulent, PDIMs/PGLs-producers and the on average less virulent, PDIMs/PGLs non-producers. In a later study it was shown that the PDIM-defect in BCG Moreau was due to a deletion in the FadD26 and the directly downstream *ppsA* gene in the same operon, which is also involved in the PDIM/PGL lipid biosynthesis pathway [39]. The authors reported that this locus was intact in BCG Japan and Glaxo. Naka *et al* later showed that BCG Tokyo 172 is actually divided into two subpopulations, in which type II, but not type I, has a frameshift mutation in the *ppsA* gene and thus does not produce PDIM lipids [40]. It is unclear whether this strain is the same as the Japan strain that was earlier reported to lack PDIMs [38]. As the Japan/Tokyo, Moreau and Glaxo substrains are not derived from one another [41], and as we have here detected a *fadD26* frameshift in a substrain derived from BCG Pasteur, these independent mutations in genes coding for key enzymes in the PDIM/PGL lipid biosynthesis pathway indicate that there may be a selective force in favor of losing this class of lipids during *in vitro* BCG cultivation. We also conclude that, as our strain has this background *fadD26* mutation, the *sapM*::Tn vaccine strain in our studies is in fact a *fadD26/sapM*::Tn double mutant, which is important knowledge for further development, as it shows that the *sapM* mutation improves vaccine efficacy even in this *fadD26* background, as is currently being used for safety enhancement of the MTBVAC lead clinical TB vaccine candidate. In addition to our *sapM*::Tn BCG mutant, this FadD26 mutation will almost certainly also be present in the Zmp1 BCG mutant candidate vaccine that was described in [42], since it is also derived from the *M. bovis* BCG Pasteur 1721 strain.

By comparative transcriptome profiling between WT BCG and *sapM*::Tn BCG, we observed an almost complete absence of *sapM* transcript in the mutant strain, together with a significant upregulation of *sapM’s* upstream gene *upp.* By RNAseq analysis, we further showed that the global transcriptional landscape in the *sapM*::Tn BCG mutant strain is not significantly altered compared to the WT BCG.

The most promising strategy for improved TB vaccines currently is the development of an improved attenuated live vaccine, either used alone or in combination with subunit booster vaccines [6]. Furthermore, re-vaccination with BCG in adolescence has recently shown promise as well [43]. An improved version of the BCG vaccine could thus be valuable in this context too. However, working with live vaccines encompasses potential safety issues, especially since a high incidence of HIV is found in areas where TB is endemic. A novel vaccine candidate should therefore be at least as safe as BCG. Safety of the *sapM*::Tn BCG strain was tested in immunocompromised SCID mice, in comparison with the WT BCG. Survival time upon infection of the mice was similar for both strains, at both high and low infectious doses, demonstrating comparable safety of the *sapM*::Tn BCG strain and WT BCG.

Due to the lack of natural infection-induced protection, the type of immune response that is crucial in preventing *M.tb* infection and TB, and which therefore should be induced by vaccination, remains largely unknown. At present, it is believed that a particular balance between differentially polarized adaptive immune responses is crucial to overcome a TB infection. *M. tb* as well as *M. bovis,* from which BCG derives, have evolved immunomodulatory mechanisms to perturb this balance to its own benefit [44,45]. We and others propose that an improved, live attenuated vaccine will need to be engineered to take down these immunomodulatory virulence factors from the pathogenic parental Mycobacteria from which these vaccines (incl. BCG) have been derived, and we propose that the SapM secreted phosphatase is such a factor. Our results demonstrate a better innate control of *sapM*::Tn BCG vaccine bacteria as compared to WT BCG upon vaccination, correlating with a faster influx of iDCs in lymphoid organs draining the vaccination site, and more modest primary expansion of TB-antigen-specific IFNγ-producing CD4^+^ and CD8^+^ T cells. Both WT BCG and *sapM*::Tn BCG have equivalent growth characteristics during *in vitro* cultivation (Suppl. Fig. 6), excluding that mere growth rate differences could have caused any difference in abovementioned phenotypes. Since complementation of *sapM* expression abolished the improved growth control of the *sapM*::Tn BCG mutant upon infection, the observed phenotypes are specifically due to the reduced expression of *sapM.* To conclude, all of this shows that the *sapM* mutated BCG vaccine behaves like a more immediately effectively controlled bacterial infection as compared to WT BCG. Importantly, in the context of viral infections, it has been well established that a protracted fight of the immune system to a chronic infection yields poorly efficacious memory due to poor generation of central memory T cells [46]. Possibly, the same holds true for chronic bacterial intracellular infections, of which BCG vaccination is an example. Together with the comparable safety of the SapM-mutated BCG and WT BCG in immunodeficient mice, and the fact that the *sapM*::Tn BCG vaccine is presently one of the few priming vaccines for which enhanced long-term survival of TB-infected animals has been demonstrated [13], these results warrant inclusion of the SapM inactivation strategy in future generations of live attenuated TB vaccines.

## 4. Materials and Methods

### 4.4 Mycobacterial strains and media

The streptomycin resistant *M. bovis* BCG Pasteur strain 1721 [26] *(RpsL,* K43R; a gift of Dr. P. Sander, Institute for Medical Microbiology, Zürich) and its *sapM* transposon insertion mutant [13] were grown in shaking culture flasks in Middlebrook 7H9 broth (Difco) supplemented with 0.05% Tween80 and Middlebrook OADC (Becton Dickinson) when grown in liquid culture. Difco Middlebrook 7H10 agar was used for growth on solid culture. For the SDS-PAGE and western blotting experiment, *M. bovis* BCG cultures were grown in 7H9 medium without OADC, but supplemented with 0.2% glucose, 0.2% glycerol and 0.05% of Tween80, because the large quantity of albumin in the OADC supplement interferes with protein electrophoresis. All strains were frozen as 1 mL cultures at OD_600_ 1 in 20% glycerol (final concentration). For all *in vitro* and *in vivo* infections described in this paper, cells were started from a glycerol stock, grown in shaking culture flasks (OD_600_ always ≤1) as described above, subcultured once and further grown until OD_600_~0.8 after which cells were collected and prepared for infection. For each experiment, we confirmed that we immunized with the same number of viable CFU by CFU plating of the inoculum.

### 4.2 Whole genome Illumina resequencing and data analysis

Genomic DNA of the *M. bovis* BCG strain 1721 and its *sapM*::Tn BCG mutant (Tn-7432) was prepared [47] and used for an Illumina library prep (Nextera XT DNA Library Preparation kit). The libraries were sequenced on an Illumina MiSeq instrument (2 × 150 bp reads), with an average 80x coverage per genome. The libraryprep and sequencing was performed by the VIB Nucleomics Core facility (www.nucleomics.be). The data was analyzed by the VIB Nucleomics Core facility and the VIB Bioinformatics Core facility (www.bits.vib.be) using CLC Genomics Workbench (CLC-GWB [48], http://www.clcbio.com/products/clc-genomics-workbench/). All reads were processed using the standard CLC-GWB settings. The reads from the *sapM*::Tn BCG strain were used for de-novo assembly in order to locate the inserted transposon. To this end, the phiMycoMarT7 transposon sequence (GenBank AF411123.1) was blasted against all de-novo contigs (n = 201). Using this information, a modified reference sequence was built from the *M. bovis* BCG Pasteur str. 1173P2 reference (NCBI NC_008769.1) by introducing the phiMycoMar T7 transposon sequence. All reads from both strains were then mapped to their corresponding reference sequences using the CLC-GWB default command settings (mismatch cost 2, insertion cost 3, deletion cost 3, length fraction 0.5, similarity fraction 0.8). A probabilistic variant analysis as implemented in the CLC-Genomics Workbench package [48] was then performed (min coverage 10, 90% variant probability) to obtain the variants listed in Table 1.

To confirm these variants, PCR primers were designed flanking the mutations, after which the regions were amplified (Phusion polymerase), purified (AMPure XP beads) and Sanger sequenced (VIB Genetics Service Facility (http://www.vibgeneticservicefacility.be) with nested primers (see table below). PCR conditions were as follows: 98°C for 3 minutes; 25 cycles of denaturation (98°C for 20 seconds), annealing (69°C for 20 seconds) and extension (72°C for 1 minute); 72°C for 5 minutes. The primer details are given in Suppl. Table 2.

### 4.3 RT-qPCR analysis

10 mL of *M. bovis* BCG cultures (grown in standard 7H9 medium until an OD_600_ of 0.8 – 1.0) were centrifuged and the pellets were washed once with 0.5% Tween80 solution in PBS. The pellet was then resuspended in 500 µl of RLT buffer (RNeasy Mini Kit, Qiagen; supplemented with β-ME). The cells were disrupted with glass beads in Retsch MM2000 bead beater at 4°C in screw-cap tubes (pre-baked at 150°C). After centrifugation (2 min, 13,000 rpm, 4°C), the supernatant was transferred to a fresh eppendorf tube. To recover the lysate trapped in between the beads, 800 µl of chloroform was added to the beads and centrifuged, after which the upper phase was transferred to the same eppendorf tube as before. Then, 1 volume of Acid Phenol/Chloroform (Ambion) was added, incubated for 2 minutes and centrifuged (5 min, 13,000 rpm, 4°C). The upper aqueous phase was transferred to a fresh eppendorf tube and this last step was repeated once. An equal volume of 70% ethanol was added and the sample was transferred to an RNeasy spin column (RNeasy Mini Kit, Qiagen). The kit manufacturer’s instructions were followed to purify the RNA. After elution in RNase-free water (30 µl), an extra DNase digestion was performed with DNaseI (10 U of enzyme, 50 µl reaction volume). Then an extra clean-up step was performed with the RNeasy Mini Kit (Qiagen). Finally, the RNA concentration was determined on a Nanodrop instrument. Typically, RNA yields of 2-3 µg were obtained with this method.

cDNA was prepared from 1 µg of DNase-treated RNA using the iScript Synthesis Kit (BioRad) and a control reaction lacking reverse transcriptase was included for each sample. The RT-PCR program was as follows: 10 minutes at 25°C, 30 minutes at 42°C, 5 minutes at 85°C, and cooling down to 12°C.

Real time quantitative PCR was done on a LightCycler 480 (Roche Diagnostics) using the SensiFast SYBR-NoRox kit (BioLine), in triplicate for each cDNA sample, on a 384-multiwell plate, with 1 ng of cDNA in a total volume of 10 µL. Primers were used at a final concentration of 10 µM. All primer pairs were generated using Primer3Plus (http://www.bioinformatics.nl/cgi-bin/primer3plus/primer3plus.cgi) as described [49]. All gene expression values were normalized using the geometric mean of the GroEL gene and 16S rRNA. Determination of amplification efficiencies and conversion of raw Cq values to normalized relative quantities (NRQ) was performed using the qbasePLUS software (Biogazelle). Statistical analysis of the NRQs was done with the Prism 6 software package using a two-tailed t-test. Primer details are given in Suppl. Table 2.

### 4.4 RNA-Seq analysis

*M. bovis* BCG cultures (*sapM*::Tn BCG mutant and WT BCG 1721) were grown in standard 7H9 medium in triplicates until an OD_600_ of 0.8 – 1.0 and RNA was prepared as described for the RT-qPCR analysis. This RNA was then depleted of ribosomal RNA (Ribo-Zero rRNA removal kit for gram-positive bacteria) and the remaining RNA was prepared for Illumina sequencing (TruSeq stranded total RNA preparation kit). This library preparation takes into account the strandedness of the RNA and is able to distinguish between sense and antisense transcripts. The libraries were sequenced on an Illumina NextSeq500 (75 bp single-end). The library preparation and sequencing was performed by the VIB Nucleomics Core facility (www.nucleomics.be). The data was analyzed using the standard RNA-seq analysis workflow in CLC Genomics Workbench v7 (CLC-GWB [48], http://www.clcbio.com/products/clc-genomics-workbench/). In short, raw sequencing reads were quality trimmed and mapped to the reference genome (*M. bovis* BCG Pasteur str. 1173P2), taking into account the read directionality. The total number of unique reads per gene (averaged over the triplicate samples) of both the WT BCG and the *sapM*::Tn BCG strain were then compared using the EdgeR statistical test [50,51].

### 4.5 Recombinant SapM production and anti-SapM antibody preparation

To express SapM in *E. coli,* we constructed an expression vector pLSPelB-SapM mature-His, being pLSAH36 carrying a PelB signal sequence, mature SapM without signal sequence and a C-terminal His tag. This was transformed to the *E. Coli* BL21+pICA2 expression strain. *E. coli* was grown in LB medium supplemented with ampicillin and kanamycin at 28°C. Expression was induced with 1 mM IPTG overnight (ON) at 18°C. Bacteria were centrifuged and the pellet was sonicated in sonication buffer (50 mM Tris, 100 mM NaCl, 1 mM PMSF, pH8). After centrifugation at 18000 rpm for 1h, supernatant was removed and inclusion bodies were extracted in extraction buffer (20 mM NaH_2_PO_4_, 0,5 M NaCl, 5 mM β-mercaptoethanol, 6 M guanidine hydrochloride, 5 mM imidazole, pH 7.5) ON at 4°C. Denatured protein was purified using nickel sepharose and eluted in elution buffer (20 mM NaH_2_PO_4_, 50 mM NaCl, 5 mM β-mercaptoethanol, 8 M guanidine hydrochloride, 400 mM imidazole, pH 7.5) and a final desalting was performed on a sephadex G25. The purification was performed by the VIB Protein Core facility. The rabbit polyclonal anti-SapM antibody was generated by immunization of 2 rabbits (CER groupe, Belgium) with purified SapM protein (625µg/mL, in the text named anti-SapM; denatured, full-length protein produced in *E. coli* as described above). Immunizations and bleedings were done following the standard CER protocol. In brief, rabbits were subcutaneously injected on days 0, 14, 28 and 56 with 500µL antigen/500µL adjuvant. Bleedings were performed on day 0(preimmune 2mL), day 38 (small test sample 2mL), day 66 (large bleed 20+2mL) and day 80 (final bleeding). Sample from the final bleeding of rabbit 2 is used in the experiments described here. The rabbit polyclonal anti-SapM-C-term antibody (Anti-PepC) was raised against a synthetic C-terminal peptide (peptide synthesis (CYATNAPPITDIWGD, terminal modification: amidation (C-terminal) and conjugation to KLH) and immunizations performed by GenScript). Immunizations and bleedings were done following the standard GenScript protocol. In brief, 2 rabbits were subcutaneously injected on days 0 and day 14. First bleedings were performed on day 0 (pre-immune, 2mL) and 21 (1mL). A third immunization was performed on day 35. A second bleeding was done on day 42 (1mL). A fourth immunization was done on day 56. Final bleeding was done on day 63. Only pre-immune serum and final anti-sera was provided by Genscript.

### 4.6 SDS-PAGE and western blotting

*M. bovis* BCG cultures were grown in 7H9 medium without OADC (as the albumin in the OADC supplement would otherwise interfere on SDS-PAGE), but supplemented with 0.2% glucose, 0.2% glycerol and 0.05% of Tween80. The supernatant samples were precipitated by first adding sodium deoxycholate (DOC, 0.1% final concentration), incubating 15 min on ice and then adding trichloroacetic acid (TCA, 7.7% final concentration) followed by 30 min of incubation on ice. The samples were centrifuged and the pellet was washed twice with acetone and dissolved again in PBS with Laemmli loading dye. Precipitated protein samples corresponding to 200 µl of culture medium were analyzed by SDS-PAGE and subsequent western blotting. The membranes were blocked overnight (4°C) with PBST (PBS + 0.05% Tween-20) containing 5% milk powder. To visualize the SapM protein, the blots were incubated for 1h at room temperature with a rabbit anti-SapM polyclonal antibody (1:5,000 dilution in PBST), washed 3 times with PBST and then incubated for 1h with a secondary goat anti-rabbit DyLight 800 conjugated antibody (30 ng/ml in PBST). After the final washing steps, the membranes were scanned with a LI-COR Odyssey system.

### 4.7 ELISA

*M. bovis* BCG culture supernatant samples (grown in 7H9; 50 µl) or control samples (0.25 µg/ml of purified SapM protein) were diluted 1:2 in 100 mM of carbonate buffer (3.03 g of Na_2_CO_3_ and 6 g of NaHCO_3_ in 1 L of water; pH 9.6) and coated on 96-well maxisorp plates (Nunc) (overnight at 4°C). The plate was then washed three times with PBST (PBS + 0.05% Tween-20). The plate was blocked with 1% of BSA in PBS (2 hours at room temperature (RT)). After plate washing, the rabbit polyclonal anti-SapM or anti-PepC antibody was added (diluted 1:5000 in PBS with 0.1% Tween-20 and 0.1% goat serum) (1 hour at RT). After washing, the plate was incubated with the secondary HRP-labeled donkey anti-rabbit antibody (GE healthcare) (diluted 1:5000 in PBS with 1% BSA) (1 hour at RT). After washing, 100 µl of the TMB substrate was added to the plate (Becton Dickinson), incubated at RT for 30 minutes, after which 50 µl of 2N H2SO4 was added to stop the reaction. The optical density at 450 nm was then determined with a bio-rad plate reader (wavelength correction at 655 nm).

### 4.8 Phosphatase assay

A 150 µl reaction mixture, containing 50 µl of *M. bovis* BCG culture supernatant samples, 85 µl of sodium acetate (0.2 M; pH 7.0) and 15 µl of para-nitrophenylphosphate (pNPP, 50 mM), was incubated for 6 hours or overnight at 37°C. Finally, 50 µl of stop buffer (1 M of Na2CO3) was added and the optical density at 415 nm was determined with a bio-rad plate reader.

### 4.9 Laboratory animals

Female C57BL/6J mice (Janvier), C57BL/6J x Balb/c mice (F1, Harlan) and CB-17 SCID mice (Charles River) were housed under specific pathogen-free conditions in micro-isolator units. At the beginning of the experiment, the mice were 7–8 weeks old. All experiments were approved by and performed according to the guidelines of the ethical committee of Ghent University, Belgium.

### 4.10 SCID safety study

Eight-week-old female CB-17 SCID mice (Charles River Laboratories, 7 or 8 mice/group as indicated) were immunized intravenously via tail vein injection with 100 µL (3 × 10^6^ versus 3 × 10^7^ cfu) of WT BCG Pasteur or *sapM*::Tn BCG mutant (diluted in PBS). The control group received PBS (pH 7.5). The animals were observed and weighed initially on a ~2-weekly basis and from day 82 post-infection on, ~every 2 days (until day 201post-infection). The differences of time to survival between the different groups were compared using log-rank test. Weight loss of 20% of initial body weight was set as the ethical endpoint.

### 4.11 Cell culture and *in vitro* infection

Bone marrow cells from C57BL/6J mice were harvested and differentiated to macrophages in DMEM containing 20% of L929-cell supernatant, 10% of heat-inactivated FCS, 0.4 mM sodiumpyruvate, 50 µM β-mercaptoethanol and 0.1 mM MEM non-essential amino acids for 7 days. Cells were cultured in antibiotic-free medium at all times. To analyze the survival/replication of *M. bovis* BCG following infection of macrophages, bone marrow-derived macrophages (BM-DMs, differentiated as described in [52]) were either left untreated or infected with WT BCG versus *sapM*::Tn BCG (MOI 10:1) in infection medium (DMEM + supplements described earlier) for 4 hours. Cells were then washed 2x with pre-warmed PBS and 1x with prewarmed medium to remove extracellular bacteria. Fresh medium was added and cells were further incubated for the indicated time points. BM-DMs were lysed in cell lysis buffer (0.05% v/v SDS, 200mM NaCl, 10mM Tris-HCl pH7.5, 5mM EDTA, 10% glycerol) at the indicated times and serial dilutions were plated onto Middlebrook 7H10 agar plates followed by incubation at 37°C.

### 4.12 Vaccination with M. *bovis* BCG and analysis of *in vivo* BCG CFU dynamics

Three experiments were performed. At the age of 8 weeks, female C57BL/6J x Balb/c (F1, Harlan) mice were infected s.c. (at the base of the tail) with 2 × 10^6^ CFU of WT BCG or *sapM*::Tn BCG mutant in 100 µl of PBS. At day 8, 14 and 28 post-infection, 10 mice/group were sacrificed by cervical dislocation; inguinal and brachial LNs were removed aseptically and homogenized in PBS-0.5% Tween 20 + EDTA-free complete protease inhibitor (Roche). Neat, 1/10, 1/100 and 1/1000 dilutions were made of which 50µL was plated in duplicate on 7H10-OADC agar and the colonies were counted after 21 days. The CFU counts were calculated from the highest dilution showing distinct colonies.

In another experiment, at the age of 8 weeks, female C57BL/6J x Balb/c (F1, Harlan) mice were infected intravenously (i.v.) with 2 × 10^6^ CFU WT BCG or *sapM*::Tn BCG mutant in 200 µl of PBS. At 24h and 3 weeks post-infection, 8 mice/group were sacrificed; spleens and lungs were removed aseptically, homogenized in PBS-0.5% Tween 20 + EDTA-free complete protease inhibitor (Roche) and analyzed for CFU counts as above. Again, neat, 1/10, 1/100 and 1/1000 (lung) and neat, 1/10, 1/100, 1/1000, 1/10 000, 1/100 000 (spleen) dilutions were made of which 50µL was plated in duplicate on 7H10-OADC agar. Additionally, 6h, 24h and 48h post-infection, blood was taken retro-orbitally and collected in BD microtainer SST tubes for serum preparation, which was frozen at -20°C until assay. Serum cytokines were determined by bioplex (BioPlex Pro Cytokine Express Assay, Biorad).

In at third experiment, at the age of 8 weeks, female C57BL/6J (Janvier) mice were infected s.c. (at the base of the tail) with 2 × 10^6^ CFU WT BCG or *sapM*::Tn BCG mutant in 100 µl PBS. At day7, 14 and 28 post-infection, 6 mice/group were sacrificed; inguinal and brachial LNs were removed aseptically and analyzed for CFU counts as above.

### 4.13 Complementation *sapM*::Tn BCG mutant

To complement the *sapM*::Tn BCG mutant strain with *sapM* gene, we constructed an integration plasmid carrying *sapM* as follows: the SapM operon was amplified by PCR using primers SapMPo-XbaI (CACTCTAGACAGTGGGTGGTCAACGACA) and Rv3311HindIII (GTTAAGCTTCCGGCTGACGGTGCCTATTC) from gDNA of BCG Pasteur. The PCR fragment was cloned in pMV261Hyg (kindly provided by Prof. E. Rubin) via XbaI and HindIII to form pMV261HygSapMoperon. The vector pMV261HygSapMoperon was digested with XbaI and ApaLI and the integrative vector pMV306Kan (kindly provided by Prof. W.R. Jacobs jr.) was digested with NheI and ApaLI, after which the appropriate fragments were purified from gel and ligated together to form pMV306HygSapMoperon, an integrating vector with the SapM operon (Plasmid map in Suppl. Fig. 7). The plasmid pMV306HygSapMoperon was electroporated into the *sapM*::Tn BCG mutant strain [53]. Hygromycin resistant colonies were selected and surviving clones were tested on a phosphatase assay.

### 4.14 Vaccination with M. *bovis* BCG and analysis of DC phenotype

C57BL/6J mice were immunized s.c. at the base of the tail with WT BCG or *sapM*::Tn BCG mutant, either with a high dose (2 × 10^6^ cfu, 5-10 mice/group) or a low dose (2 × 10^5^ cfu, 10 mice/group). Seven and 14 days later, the mice were sacrificed, and inguinal LNs, brachial LNs and spleens were isolated. Cells were prepared by crushing the organs on a 70µm cell strainer and labelled with an optimized DC-analysis antibody panel and analyzed by flow cytometry. Fluorochrome-conjugated mAbs specific for mouse CD11c (clone N418, ebioscience), CD8 (clone 53-6.7, BD), CD40 (clone 1C10, ebioscience), CD80 (clone 16-10A1, Biolegend), Ly-6C (clone AL-21, BD), MHCII (clone M5/114.15.2, ebioscience), CD103 (clone 2E7, ebiosciene), F4/80 (clone BM8, ebioscience), CD11b (clone M1/70.15, Life Technologies), Ly-6G (clone RB6-8C5, ebioscience) were used. Dilutions of the antibodies are given in Suppl. Table 3 and gating strategy is given in Suppl. Fig. 8.

### 4.15 Vaccination with M. *bovis* BCG for T cell phenotyping and antigen-specific cytokine determination

C57BL/6J mice were vaccinated s.c. at the base of the tail with either a high dose (2 × 10^6^ cfu) or a low dose (2 × 10^5^ cfu, in 100µL of PBS) WT BCG or *sapM*::Tn BCG mutant. 14 days after infection, mice were killed by cervical dislocation and inguinal + brachial LNs and spleens were removed aseptically. Cells were prepared by crushing the organs on a 70 µm cell strainer and adjusted to a concentration of 1 × 10^6^ (LN) or 2 × 10^6^ (spleen) cells/ mL in either flat-bottom 96-well microwell plates (Bioplex/ELISA) or round-bottom plates (flow cytometry) (Nunc), in RPMI 1640 medium, supplemented with 10% fetal calf serum, L-glutamine (0.03%), 0,4 mM sodiumpyruvate, 0,1 mM non-essential amino acids and 50 µM β-mercaptoethanol. Cells were restimulated with purified native Ag85A (Rv3804c) from *M. tb,* strain H37Rv (15 µg/mL in PBS, obtained through BEI Resources, NIAID, NIH), purified protein derivative (PPD, 10 µg/mL, produced at Scientific Institute for Public Health, Brussels, Belgium) and anti-CD3/28 (10 µg/mL, BD) in a volume of 100 µL added to 100 µL of cell suspension. For surface and intracellular staining, cells were incubated at 37°C in a humidified CO2 incubator, and after 1h, brefeldin was added (10 µg/mL-Sigma Aldrich) for another 5 hours. LNs and splenocytes were subsequently labelled with an optimized antibody panel for T cell phenotyping and analyzed by flow cytometry. Fluorochrome-conjugated mAbs specific for mouse CD3 (clone 17A2, BD), NK1.1 (clone PK136, BD), CD4 (clone RM4-5, Invitrogen), CD8 (clone 53-6.7, BD), CD335 (clone 29A1.4, BD), TCRyô (clone GL-3, ebioscience), IL17 (clone TC11-18H10, BD), IFNγ (clone XMG1.2, BD) and TNF (clone MP6-XT22, ebioscience) were used. Fixable viability dye eFluor 780 (eBioscience) was used as a live/dead marker.

Dilutions of the antibodies and gating strategy are given in Suppl. Materials and Methods. For determination of cytokine levels in the supernatant, the latter was harvested after 48h post-stimulation. Supernatants were frozen at -20°C until assay. Cytokines were determined by bioplex (BioPlex Pro Cytokine Express Assay, Biorad) and ELISA (mouse IFNγ duoset Elisa, R&D systems) as indicated in the text.

### 4.16 Flow cytometry analysis

Cells were first blocked by adding Fc block (purified rat anti-mouse CD16/CD32 monoclonal Ab, clone 2.4G2, BD, 0.1µg/sample) and incubating for 15 min at 4°C. Staining is performed for 20 min at 4 °C in the dark. For intracellular staining, cells were incubated for 5 h with 10 µg/mL brefeldin A (10 µg/mL-Sigma Aldrich) before fixation and permeabilization (Fixation-Permeabilization concentrate, Permeabilization buffer, ebioscience). In all analyses, following doublet exclusion, live cells were identified using a fixable viability dye (Molecular Probe, Life Technologies). Data were acquired on an LSR II equipped with three lasers (488, 635, 405nm) (APC panel) or Fortessa with 4 lasers (405, 488, 561, 633 nm) (T cell panel) (BD Biosciences) and analyzed using FlowJo software (Tree Star, Ashland, OR).

### 4.17 Statistical analysis

Results are presented as means ± standard error of the mean (SEM) unless otherwise stated and groups were compared using statistical tests as mentioned in the text, using Prism Software (GraphPad Software, San Diego, CA).

## Supporting information

## 5. Acknowledgements

This work was supported by an ERC Consolidator grant to N.C. (GlycoTarget; 616966) and a GOA grant. N.F. and D.V. were postdoctoral fellows and K.V. and K.B. were predoctoral fellows of FWO (Fonds Wetenschappelijk Onderzoek-Vlaanderen). We thank Dr. Peter Sander (Institute for Medical Microbiology, Faculty of Medicine, University of Zurich) for providing us with the *M. bovis* BCG strain (strain 1721, *RpsL,* K43R), Dr. S. Alonso (Dept, Microbiology, National University of Singapore) for providing us the optimal protocol for electroporation of *M. bovis* BCG and Dr. K. Huygen (WIV, Brussels) for providing us with the PPD.

We thank Prof. Dr. E. Rubin (Dept, Immunology and Infectious diseases, Harvard School of Public Health, Boston) for providing us the pMV261Hyg vector and Prof. W.R. Jacobs Jr (Albert Einstein College of Medicine, New York) for providing us the pMV306Kan vector. We thank the VIB Nucleomics core (www.nucleomics.be) and Genomics Core (UZ – K.U. Leuven, gc.uzleuven.be) facilities for the Illumina MiSeq and HiSeq sequencing services. We thank Jannick Leoen from the VIB Protein Core facility (https://corefacilities.vib.be/psf) for purification of SapM protein. The following reagent was obtained through BEI Resources, NIAID, NIH: Ag85 Complex, Purified Native Protein from *Mycobacterium tuberculosis,* Strain H37Rv, NR-14855.

N.C and N.F. are named inventors on a patent covering the use of SapM mutation as a vaccine improvement strategy.

## 6. Author contributions

NF coordinated the project, designed and performed experiments, analyzed and interpreted data and wrote the manuscript. KV analyzed genome sequencing and RNAseq experiments, performed RT-qPCR experiments and co-wrote the manuscript. EH designed and performed *in vivo* experiments, analyzed and interpreted data. EP performed all *in vitro* experiments and helped in performing *in vivo* experiments. DV and KB helped in performing *in vivo* experiments. PT and AV made the *sapM*::Tn:complementation mutant. NC coordinated the project, interpreted data and co-wrote the manuscript.

