## Supplementary material for "SapM mutation to improve the BCG vaccine: genomic, transcriptomic and preclinical safety characterization"

### Supplementary Figures

**Supplementary Figure 1| Whole genome resequencing of the WT BCG Pasteur 1721 strain and of the *sapM::Tn* BCG mutant.** Graphical representation of the rather uniform coverage obtained after resequencing the genome of both the WT BCG and *sapM::Tn* BCG mutant strains. Average coverage of ± 80x.

**Supplementary Figure 2| (A) Correlation dot plot of the averaged unique gene counts between WT BCG and *sapM*::Tn BCG samples.** The diagonal line is set at x – y. **(B) Correlation dot plots of the unique gene counts between the individual WT BCG and *sapM*::Tn BCG replicates.** For clarity, only the lower left fraction of the dot plot is depicted (black square in Suppl. Fig. 2A). The *sapM*, *upp* and *BCG_3376* genes are shown in red. The diagonal line is set at x – y.

**Supplementary Figure 3| *In vivo* replication analysis**

C57BL/6J mice were vaccinated i.v. with the parental (black) or *sapM*::Tn *M. bovis* BCG (red) ($2.10^6$ cfu, 8 mice/group). **(A)** At 24h and 3 weeks post-infection, mice were sacrificed and the number of bacteria in the spleens was determined by cfu plating (Mann-Whitney U test; ** P<0.01). **(B)** F1 mice were vaccinated i.v. with the parental or *sapM*::Tn *M. bovis* BCG ($2.10^6$ cfu, 8 mice/group). 6h, 24h, 48h post-infection, mice were bled and cytokines were determined in the serum by bioplex (Mann-Whitney test; ** P<0.01 *P<0.05).

**Supplementary Figure 4| Vaccination with *sapM*::Tn BCG induces reduced frequencies of IFNγ-producing $CD4^+$ and $CD8^+$ T cells compared to the parental strain**

C57BL/6J mice were immunized s.c. at the base of the tail with a low dose of *M. bovis* BCG WT (black) or *sapM*::Tn mutant (red) ($2.10^5$ cfu, 10 mice/group). 14 days later, the mice were sacrificed, and inguinal LNs, brachial LNs and spleens were isolated. Cells were prepared and the T cell response was analyzed by intracellular cytokine staining followed by flow cytometry. (Mann-Whitney or 2way ANOVA; *P<0.05, **P<0.01)

**Supplementary Figure 5| Reduced frequencies of IFNγ-producing $CD4^+$ and $CD8^+$ T cells compared to the parental strain upon vaccination with *sapM*::Tn BCG are absent at later time points**

C57BL/6J mice were immunized s.c. at the base of the tail with a high dose of *M. bovis* BCG WT (black) or *sapM*::Tn mutant (red) ($2.10^6$ cfu, 10 mice/group). 6 weeks later, the mice were sacrificed, and inguinal LNs, brachial LNs and spleens were isolated. Cells were prepared and the T cell response was analyzed by intracellular cytokine staining followed by flow cytometry. (Mann-Whitney or 2way ANOVA; *P<0.05, **P<0.01)

**Supplementary Figure 6| I*n vitro* growth of *M. bovis* BCG WT and *sapM::Tn* mutant**

*M. bovis* BCG WT and *sapM*::Tn BCG mutant were grown in shaking culture flasks in 7H9 broth supplemented with 0.05% Tween80 and Middlebrook OADC. On various time points, $OD_{600}$ was determined. An average of 2 independent experiments (with indication of SEM) is shown in the figure.

**Supplementary Figure 7| Plasmid map of pMV306HygSapMoperon**

**Supplementary Figure 8| Gating strategy for APC and T cell panel**

### Supplementary Tables

**Supplementary Table 1 | Excel file (CLC output) of differential expression**

The first tab contains a legend, explaining all columns and data in the second tab.

**Supplementary Table 2 | Primer tables**

Upper table: PCR primers to confirm the variants described in Table 1; Lower table: Primers that were used for RT-qPCR described in paragraph 4.3.

**Supplementary Table 3 | Dilutions of used antibodies**

### Accession codes

The novel sequencing data generated in this study has been deposited at NCBI's Sequence Read Archive under BioProject codes PRJNA506330 and PRJNA506333.
