## Supplementary material for "SapM mutation to improve the BCG vaccine: genomic, transcriptomic and preclinical safety characterization"

Suppl. Fig. 1

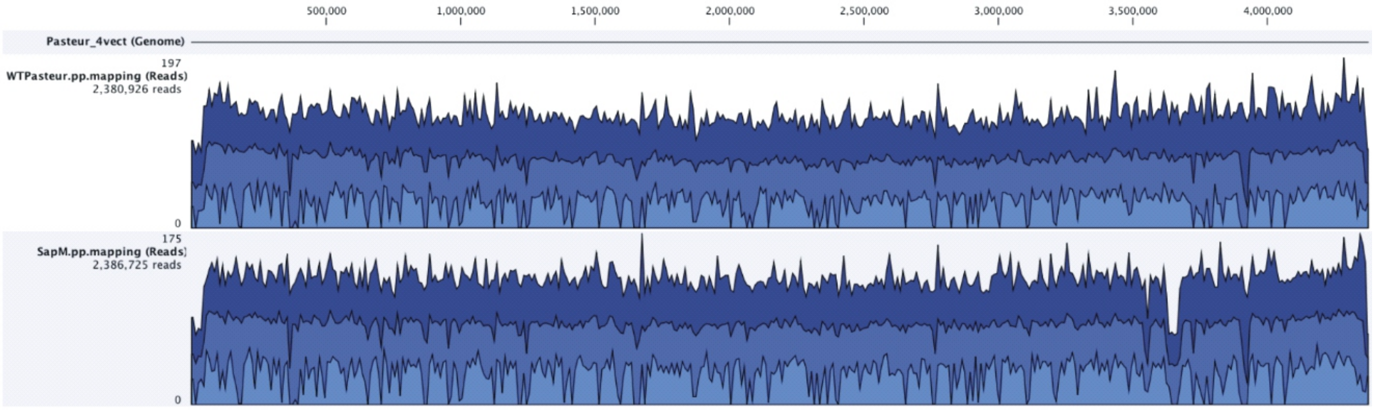
500,000
1,000,000
1,500,000
2,000,000
2,500,000
3,000,000
3,500,000
4,000,000
Pasteur_4vect (Genome)
197
WTPasteur.pp.mapping (Reads)
2,380,926 reads
0
175
SapM.pp.mapping (Reads)
2,386,725 reads
0

Suppl Fig. 2

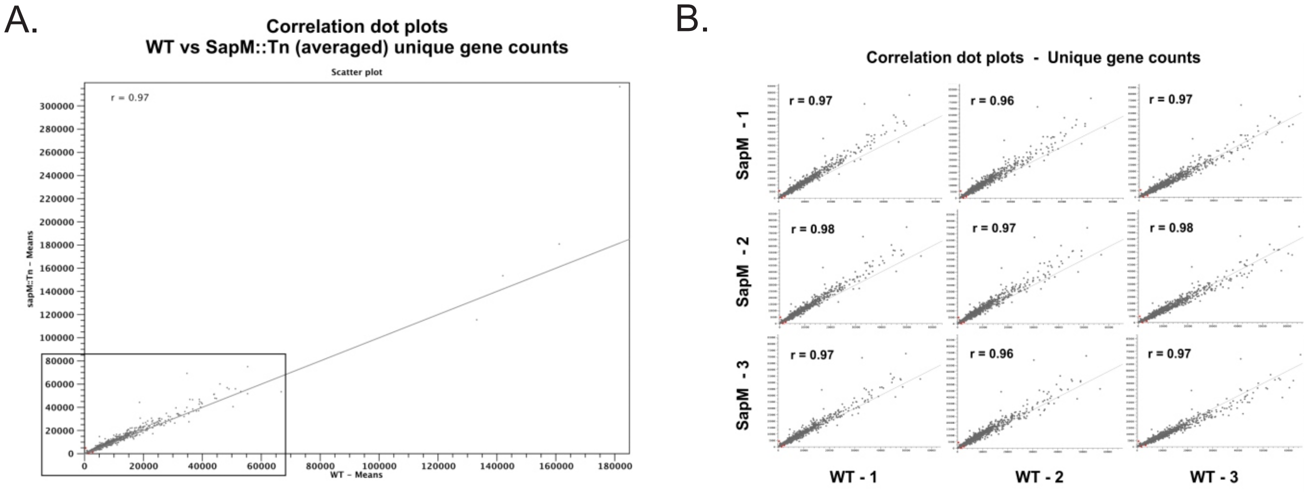
A.
Correlation dot plots
WT vs SapM::Tn (averaged) unique gene counts
Scatter plot
r = 0.97
sapM:Tn – Means
WT – Means
B.
Correlation dot plots - Unique gene counts
r = 0.97
r = 0.96
r = 0.97
r = 0.98
r = 0.97
r = 0.98
r = 0.97
r = 0.96
r = 0.97
SapM - 1
SapM - 2
SapM - 3
WT - 1
WT - 2
WT - 3

Suppl Fig. 3

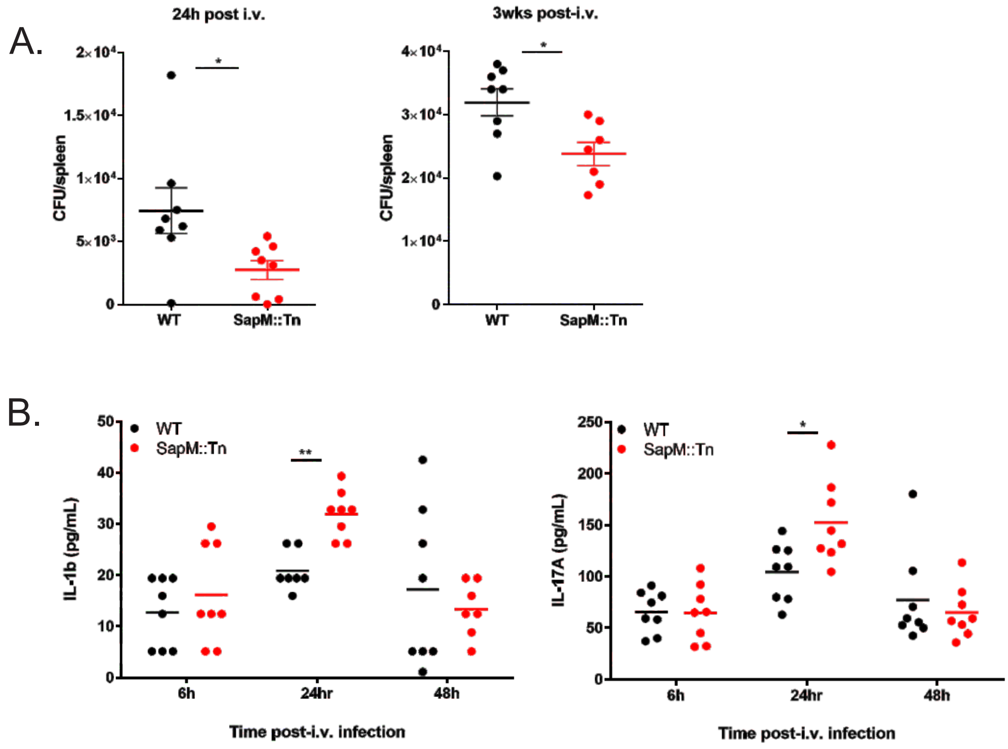
A.
24h post i.v.
3wks post-i.v.
CFU/spleen
WT
SapM::Tn
B.
WT
SapM::Tn
IL-1b (pg/mL)
IL-17A (pg/mL)
6h
24hr
48h
Time post-i.v. infection

Suppl Fig. 4

Vaccination-site draining lymph node; 2 weeks post-vaccination

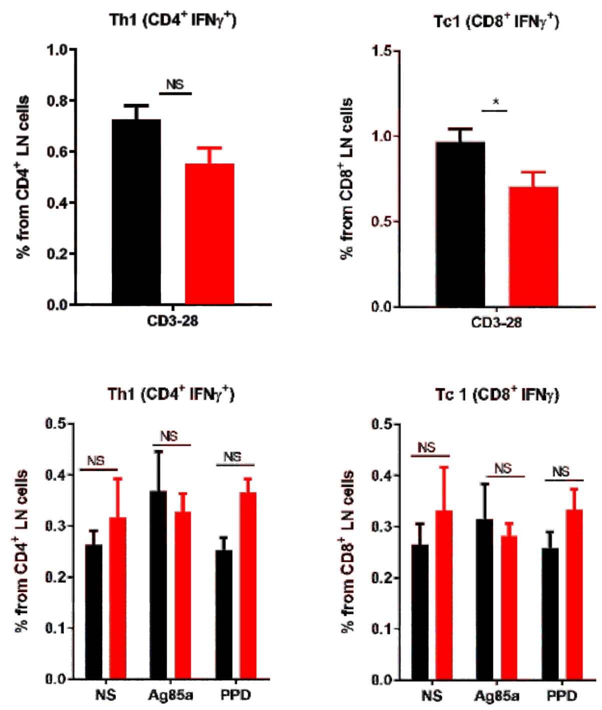


Spleen; 2 weeks post-vaccination

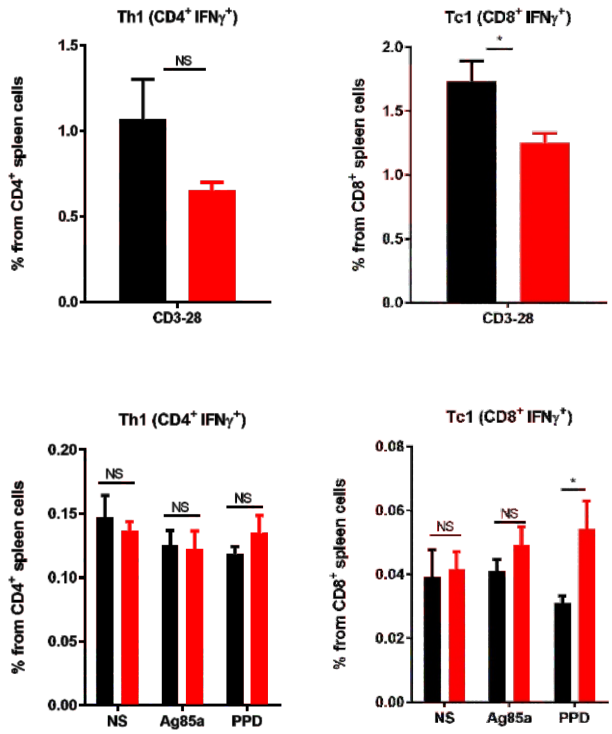

### Suppl Fig. 5

Vaccination-site draining lymph node; 6 weeks post-vaccination

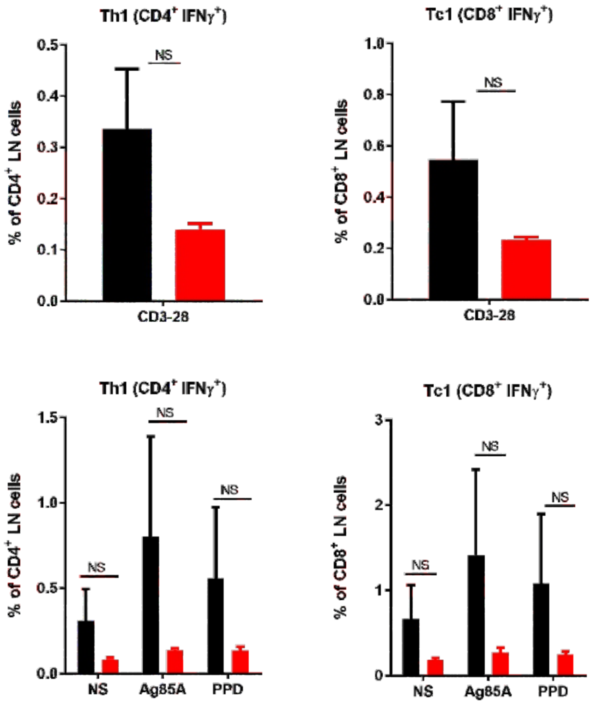


Spleen; 6 weeks post-vaccination

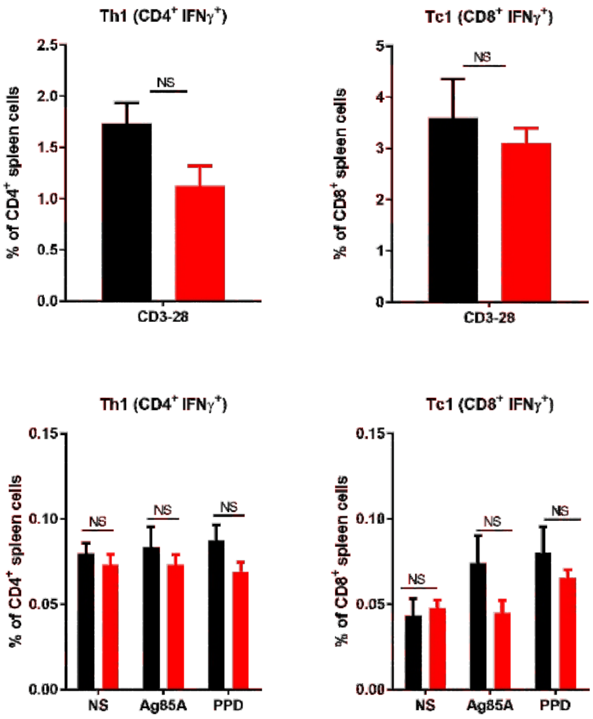

Suppl Fig. 6

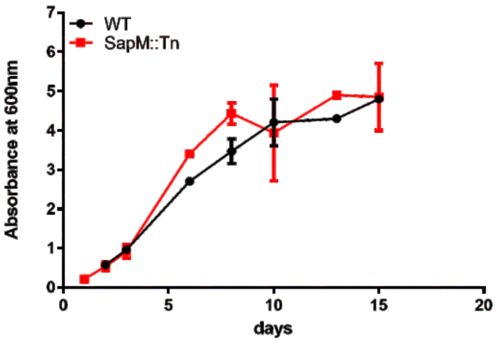

Suppl Fig. 7

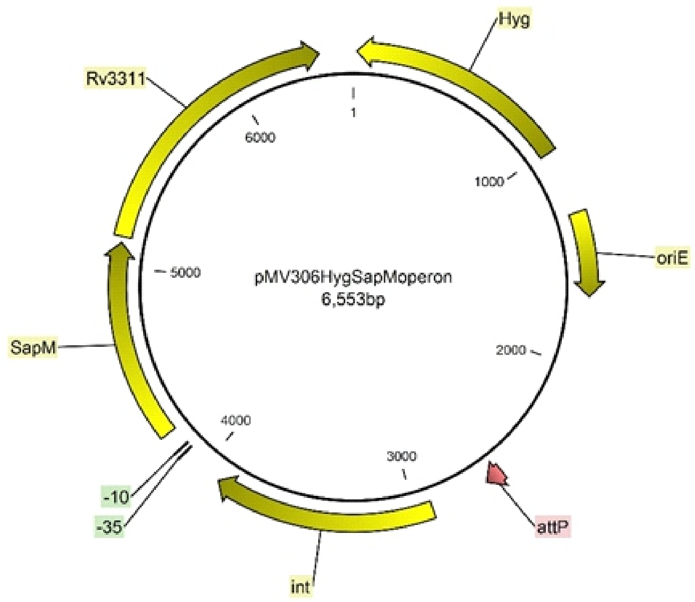
Hyg
Rv3311
1
6000
1000
oriE
5000
pMV306HygSapMoperon
6,553bp
SapM
2000
4000
3000
-10
-35
attP
int

Suppl Fig. 8

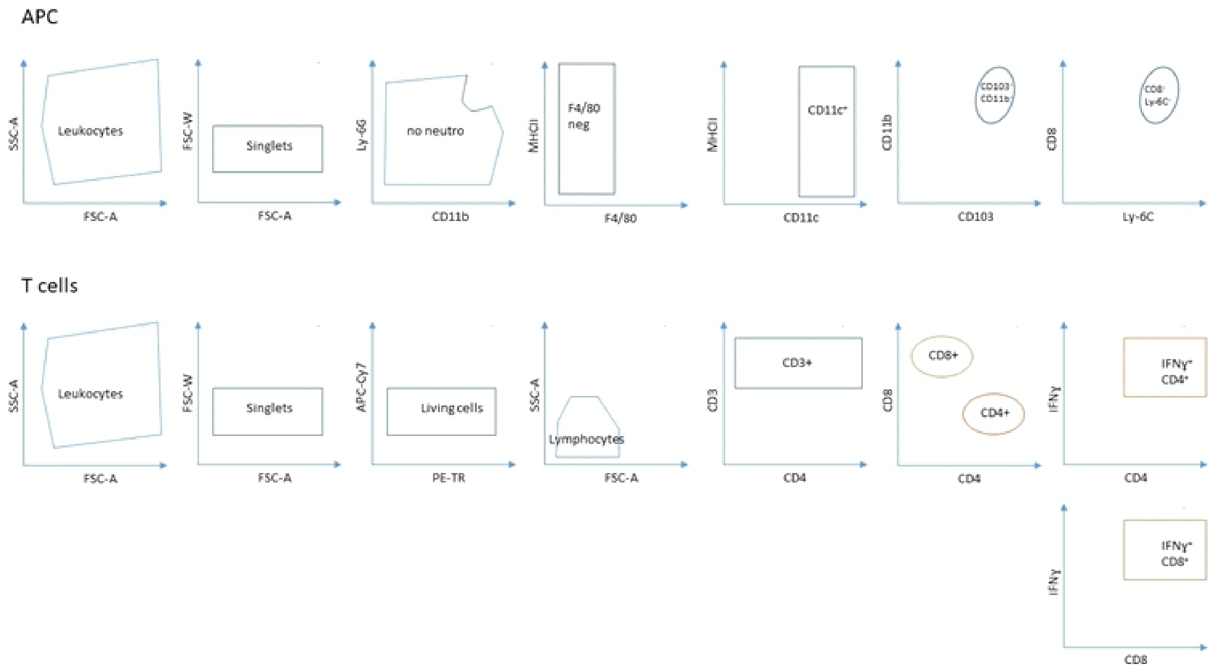
APC
SSC-A
Leukocytes
FSC-A
FSC-W
Singlets
FSC-A
Ly-6G
no neutro
CD11b
MHCII
F4/80
neg
F4/80
MHCII
CD11c
CD11c
CD11b
CD103
CD8
Ly-6C
T cells
SSC-A
Leukocytes
FSC-A
FSC-W
Singlets
FSC-A
APC-Cy7
Living cells
PE-TR
SSC-A
Lymphocytes
FSC-A
CD3
CD3+
CD4
CD8
CD8+
CD4+
CD4
IFNγ
CD4
IFNγ
CD8
