## Supplementary material for "SapM mutation to improve the BCG vaccine: genomic, transcriptomic and preclinical safety characterization"

| Explanation of data output table (WT-7H9 vs. SapM::Tn-7H9) | | |
|---|---|---|
| **Column names** | | **Explanation** |
| **Feature ID** | | Standard locus ID of each gene in the M. bovis BCG pasteur genome. |
| **Experiment (original values)** | | Output of the comparison WT-7H9 vs SapM::Tn-7H9. More information in the online manual (http://resources.qiagenbioinformatics.com/manuals/clcmainworkbench/800/index.php?manual=Experiment_level.html). |
| | **Range** | The 'Range' column contains the difference between the highest and the lowest expression value for the feature over all the samples. If a feature has the value NaN in one or more of the samples the range value is NaN. |
| | **IQR** | The 'IQR' column contains the inter-quantile range of the values for a feature across the samples, that is, the difference between the 75 %-ile value and the 25 %-ile value. For the IQR values, only the numeric values are considered when percentiles are calculated (that is, NaN and +Inf or -Inf values are ignored), and if there are fewer than four samples with numeric values for a feature, the IQR is set to be the difference between the highest and lowest of these. |
| | **Difference** | Difference of the mean expression values ("Means" column) of the SapM::Tn-7H9 condition minus the WT-7H9 condition. |
| | **Fold Change** | Ratio of the mean expression values ("Means" column) of the SapM::Tn-7H9 condition over the WT-7H9 condition. |
| **EDGE test: WT-7H9 vs SapM::Tn-7H9, tagwise dispersions** | | Output of the statistical test "Empirical analysis of DGE" as implemented in CLC Genomics Workbench. More information in the online manual (http://resources.qiagenbioinformatics.com/manuals/clcgenomicsworkbench/702/index.php?manual=Empirical_analysis_DGE.html) |
| | **P-value** | The 'P-value' holds the p-value for the Exact test in the EDGE algorithm. |
| | **Fold change** <br> **Weighted difference** | The 'Fold Change' and 'Weighted difference' columns are both calculated from the estimated 'average cpm (counts per million)' values of each of the groups. The estimated 'average cpm' values are values that are derived internally in the Exact Test algorithm. They depend on both the sizes of the samples, the magnitude of the counts and on the estimated negative binomial dispersion, so they cannot be obtained from the original counts by simple algebraic calculations. The **'Fold Change'** will tell you how many times bigger the average cpm value of group 2 is relative to that of group 1. If the average cpm value of group 2 is bigger than that of group 1 the fold change is the average cpm value of group 2 divided by that of group 1. If the average cpm value of group 2 is smaller than that of group 1 the fold change is the average cpm value of group 1 divided by that of group 2 with a negative sign. **The 'weighted difference'** column contains the difference between the average cpm value of group 2 and the average cpm value of group 1. |
| | **FDR p-value correction** | Corrected p-value using the false disovery rate (FDR) method. The false discovery rate is the proportion of false positives among all those declared positive. We expect 5 % of the features with FDR corrected p-values below 0.05 to be false positive. There are many methods for controlling the FDR - the method used in CLC Genomics Workbench is that of (Benjamini and Hochberg, 1995, Journal-Royal Statistical Society Series B). |
| **WT-7H9 / SapM::Tn-7H9** | | Two conditions (strains) that were compared in the RNA-seq experiment. Three replicate samples were analyzed per condition. |
| | **Expression on values** | Expression value per gene. Identical to the "Total gene reads" column as these were the values that were selected for comparison between conditions. |
| | **Unique gene reads** | Number of reads that uniquely map to each gene (or "Feature ID"). |
| | **Total gene reads** | Total number of reads that map to each gene (or "Feature ID"). Identical to the "Unique gene reads" column as reads that map non-uniquely were not considered during the mapping algorithm. |
| | **RPKM** | RPKM, Reads Per Kilobase of exon model per Million mapped reads (Mortazavi et al, 2008, Nature Methods). For a bacterial genome, the RPKM = (total gene reads) / (mapped reads (millions) x gene length (kb)). |
| | **Chromosome (Chr)** | Chromosome name on which the "Feature ID" tag is located. |
| | **Chr region start** | Start site (in bp) of the "Feature ID" tag on the genome. |
| | **Chr region end** | Stop site (in bp) of the "Feature ID" tag on the genome. |
| | **Means** | Average of the expression values (column "Expression on values") of the replicate samples. |

**Supplementary Table 1**
