## Supplementary material for "SapM mutation to improve the BCG vaccine: genomic, transcriptomic and preclinical safety characterization"

| Variant | gene | position SNP | PCR primers | | Sequencing primers | |
|---|---|---|---|---|---|---|
| 1 | rpsL | 813,096 | Fw | GTCATCGCCGTGCTGTCCTC | Fw | GGTACCGCGTCAACGTAGCG |
| | | | Rev | CCTTGTTCACCAACTGGGTGACC | Rev | CTTCTTAGCGCCGTAACGGCTG |
| 2 | BCG_1242c – pks3 | 1,344,671 | Fw | GTCCGACGACGGAAAACTGTTATC | Fw | CAGGCCGATCATCGTCTTGC |
| | | | Rev | CTGTACCAGCCGTGGGTGAATG | Rev | GCTGCAAGGTCCGATACAACG |
| 3 | fadD26 | 3,197,940 | Fw | CATAGTGAACGCCAGAAAGCCG | Fw | AGCAGCCTGACAGCACTGCATATAC |
| | | | Rev | ATTGGCAATGACATTCGTGTGC | Rev | GGCGAGTCCAAATCAAGCAG |
| 4 | BCG_3499c | 3,833,491 | Fw | TCACGAAGGTGGCCAGATTCAG | Fw | GGTGAACATCCGACCGGTGTAG |
| | | | Rev | TTGTGGTTCGCGCTGGACAC | Rev | CTCAAGGGCAATGTCACCGTC |
| 5 | cut3 | 3,854,971 | Fw | CATACGCAGACAACGTCGCAG | Fw | CTATTCGGTTCCAAGGCGATTG |
| | | | Rev | GCCAAGAAATCACCGTTGGC | Rev | CGCAATGAACTGACGAAAGCTTG |
| | sapM | 3,685,759 | Fw | CATCGGGTCAAGCACCATGAC | Fw | AGAGACGCTCTCGAAGCCATACAGG |
| | | | Rev | GCCCACCTGCAACCAGAAGTC | Rev | CCAGGGTATACCGTGCCTTGG |
| 6 | SugI | 3,708,425 AND 3,708,517 | Fw | GACACTGATCGAACCCGACC | Fw | TGTTCGGGTTGATCATGGTCG |
| | | | Rev | GCTAAGCGTCGCCAGCTGATAC | Rev | GATGAGCACCACCGATTTCTTG |

| gene | RT-qPCR primers | |
|---|---|---|
| 16S | Fw | ATGACGGCCTTCGGGTTGTAA |
| | Rev | CGGCTGCTGGCACGTAGTTG |
| GroEL | Fw | CGAGGCGATGGACAAGGT |
| | Rev | CTTGTCGAACCGCATACCCT |
| Upp1 | Fw | GGGCAGCGAGTCCAGATA |
| | Rev | CACGCTGCTGTTGATCTATG |
| Upp2 | Fw | CGGGTCGCGGCTAAC |
| | Rev | GGGCAGCGAGTCCAGATA |
| SapM | Fw | AGGGTATACCGTGCCTTGG |
| | Rev | GTTCTCCTCCACCACGATG |
| SapMb | Fw | TGCGGCCCGGAACTTACAACGAGA |
| | Rev | CAAGCGGATGGGTACGAGGTCAGC |
| BCG_3376_3 | Fw | ATCGACGAGATCATCAGCAC |
| | Rev | CCCAGTACGACACTGTCACC |
| BCG_3376_3 | Fw | AAGTTCTTCAACGGCAATCC |
| | Rev | GTGCTGATGATCTCGTCGAT |

Supplementary Table 2
