## Supplementary material for "SapM mutation to improve the BCG vaccine: genomic, transcriptomic and preclinical safety characterization"

**Supplementary Table 3: Dilutions of used antibodies**

**Surface panel APCs**

| | stock conc. | dilution |
|---|---|---|
| CD11c PercpCy5.5 | 0,2 mg/mL | 1:100 |
| CD8 V500 | 0.2mg/ml | 1:100 |
| CD40 APC | 0.2mg/ml | 1:100 |
| CD80 BV421 | 0.05mg/ml | 1:50 |
| Ly-6C PeCy7 | 0.2mg/ml | 1:200 |
| MHCII FITC | 0.5mg/ml | 1:400 |
| CD103-PE | 0.2mg/ml | 1:100 |
| F4/80 APC-eFluor780 | 0.2mg/ml | 1:100 |
| CD11b- PE-TR | 0.2mg/ml | 1:200 |
| Ly-6G-AF700 | 0.2mg/ml | 1:100 |

**Surface panel T cells**

| | stock conc. | dilution |
|---|---|---|
| CD3 FITC | 0,5mg/mL | 1:150 |
| NK1.1 BV605 | 0,2mg/mL | 1:100 |
| CD4 PE-TexasRed | 0,2mg/mL | 1:200 |
| CD8 V500 | 0,2mg/mL | 1:100 |
| CD335 BV421 | 0,2mg/mL | 1:50 |
| TCRγδ APC | 0,2mg/mL | 1:100 |
| L/D stain eFluor780 | | 1:200 |

**ICS**

| | | |
|---|---|---|
| IL17 PE | 0,2mg/mL | 1:50 |
| IFNγ AF700 | 0,2mg/mL | 1:50 |
| TNF PerCP-Efluor710 | 0,2mg/mL | 1:100 |
